# Generative Approaches to Kinetic Parameter Inference in Metabolic Networks via Latent Space Exploration

**DOI:** 10.1101/2025.03.31.646317

**Authors:** Subham Choudhury, Ilias Toumpe, Oussama Gabouj, Jakob Sebastian Behler, Vassily Hatzimanikatis, Ljubisa Miskovic

## Abstract

Generative machine learning methods that utilize neural networks to parameterize large-scale and near-genome-scale kinetic models have yielded significant efficiency gains in model construction, paving the way for high-throughput dynamic metabolism studies in biomedical and biotechnological applications. Nevertheless, challenges remain in interpreting the outputs of generative neural networks and developing strategies to quickly adapt these networks to different organisms and physiological contexts without having to restart the modeling process from scratch. Here, we present a systematic framework for repurposing generative neural networks trained under one physiological context to build large-scale kinetic models tailored to another. We showcase the effectiveness of this framework through three case studies in *Escherichia coli*: (i) adjusting the speed of the dynamic response of aerobic metabolism, (ii) improving interpretability by identifying key enzymatic steps that limit the dynamic response speed of the metabolic models, and (iii) adapting a trained generator to capture the distinct dynamic behavior of anaerobic metabolism. To assess robustness and generalizability beyond *E. coli*, we extend our approach to large-scale kinetic models of *Saccharomyces cerevisiae*, systematically exploring latent-space–driven control of network dynamics across generators at different training stages and across multiple representative regions of the latent input space. Together, these results demonstrate that latent space exploration provides a transferable and computationally efficient strategy for controlling the dynamic behavior in large-scale kinetic models across species and physiological regimes. Given the growing adoption of generative neural networks in biological systems modeling, our approach has the potential to facilitate applications in personalized medicine and accelerate the high-throughput design of cell factories by streamlining model construction across diverse living organisms.

## INTRODUCTION

Metabolism is an inherently adaptive dynamic process that sustains cellular functions and growth despite ongoing changes in the extracellular environment or the cell’s internal state. Accurately describing the systems-wide behavior of metabolic processes is essential for advancing biomedical and biotechnological applications, such as the development of personalized therapies^1,2^, understanding metabolic shifts between healthy and diseased states^3^, engineering cellular factories for biochemical and pharmaceutical production^4,5^, and uncovering and harnessing metabolic interactions within microbial communities^6–8^. Driven by recent advancements in omics technologies, modeling techniques, and computational tools, mathematical models play an increasingly important role in this task.^9,10^

The adaptability of metabolic processes to external and internal changes can be captured by dynamic (kinetic) models, which can depict the evolution of metabolite concentrations, metabolic fluxes, and enzyme concentrations over time. By incorporating information about enzyme kinetics and regulatory interactions, these models provide a quantitative framework to investigate metabolic responses to genetic and environmental perturbations, regulatory dynamics in metabolism, and its intricate interplay with other cellular processes. Dynamic models have demonstrated significant potential in diverse areas, including targeted drug development^11,12^, bioproduction^13–15^, environmental bioremediation^16^, and the study of multi-species microbial ecologies^17^. They are also well-suited for integrating and reconciling diverse biological datasets, including multi-omics data.^18,14,19^

However, our incomplete knowledge of key biochemical and biophysical properties, such as enzyme kinetics, thermodynamics, and regulatory interactions, makes it challenging to determine the parameters underlying fundamental dynamics and intrinsic complexities of metabolic processes.^20^ Because of the scarcity of available experimentally measured parameters^21,22^, considerable efforts have been invested in efficiently parameterizing these models using traditional approaches such as Monte Carlo sampling^23–25^, and hybrid or ensemble modeling^26,18,27^. More recently, it was understood that integrating dynamic models with Machine Learning (ML) approaches can enhance their accuracy, scalability, and robustness.^28–32,19^

Notably, generative ML-based parameterization methods, like REKINDLE^31^ and RENAISSANCE^19^, have demonstrated efficiency gains of several orders of magnitude over traditional kinetic modeling frameworks. Such generative methods employ neural networks, so-called generators, to parameterize large-scale and near-genome-scale dynamic models with desirable dynamic metabolic properties.

Once trained, these generators can be fine-tuned with a low amount of data via transfer learning^33,34^ for studying similar physiologies, cell lines, or patients. However, to bring high-throughput dynamic studies into a practical reality, several key challenges must be addressed. In particular, how can we repurpose and improve models built for one physiology or satisfying a specific set of model properties for use in studies with entirely different requirements? For example, can we repurpose a generator trained to produce models of a wild-type strain to obtain models that capture both wild-type and recombinant physiologies? Similarly, how can we use a generator initially trained to produce models for a slow-growing strain to generate models for fast-growing strains? Additionally, while improving model construction efficiency and accuracy is vital, it is also necessary to address the challenge of systematically interpreting generator outputs to extract biologically meaningful insights. For instance, can these methods provide information useful for guiding the design of experimental strategies for biotechnological applications without requiring downstream studies^14^?

A venue to address these challenges is to explore the generator’s latent space — an abstract, lower-dimensional representation of the kinetic parameter space that is explicitly learned in RENAISSANCE and REKINDLE owing to their specific generative architectures, and which is mapped during successful training of a generator for the studied physiology (Figure 1a), thereby enabling systematic post-training exploration and control of model properties. Latent space encodings of complex datasets are routinely used in various classes of machine learning algorithms^35,36^. These encodings map multidimensional, often rugged parametric spaces onto simpler, lower-dimensional latent spaces. Machine learning algorithms can produce these nonlinear mappings between the two spaces by training on large datasets from the system of interest. These mappings can then be used to sample the latent space and generate new instances from the dataset’s parametric space. Recent efforts^37–39^ have focused on shedding light on the ‘black box’ nature of these advanced ML algorithms, with significant attention given to understanding how the features (attributes) of high-dimensional data are distributed within their latent spaces. Some studies have shown that features of complex datasets can be linearly decoupled in latent space^40,41^. This decoupling has been used to guide and control the generation process through regression-based^42^ or semantic techniques^38^, allowing the generation of new data points with specific or conditional features^31,43^, or a weighted sum of features not present in the original dataset^41^. This means that the latent space of machine learning algorithms allows for the model reduction of complex systems and provides more flexible control over the system’s properties.

In this study, we present a framework that leverages the reduced size and structured landscape of the generator’s latent spaces to produce dynamic models tailored to the studied physiology (Figure 1).

The framework begins by employing state-of-the-art generative ML methods, such as REKINDLE or RENAISSANCE, to create a generator that maps a simple Gaussian latent space to the relevant sub-space of kinetic parameters constrained by integrated reference data (Figure 1b). Latent space perturbations are then nonlinearly transformed into the parametric space through the trained generator, producing novel kinetic parameter sets with distinct dynamic responses and properties (Figure 1c, Supplementary Note 1). This framework offers a systematic and efficient approach to exploring the dynamic properties of metabolic models. We illustrate its robustness and generalizability by applying latent space exploration to large-scale kinetic models of *Saccharomyces cerevisiae*. We further demonstrate its versatility through three applications in *E. coli*: (I) Fine-tuning modeled dynamic response times in aerobically grown *cells*; (II) Identifying dynamic bottlenecks at the enzyme level; and (III) repurposing trained neural network generators to create dynamic models meeting the distinct dynamic requirements of anaerobically grown *cultures* (Figure 1d).

**Figure 1.**
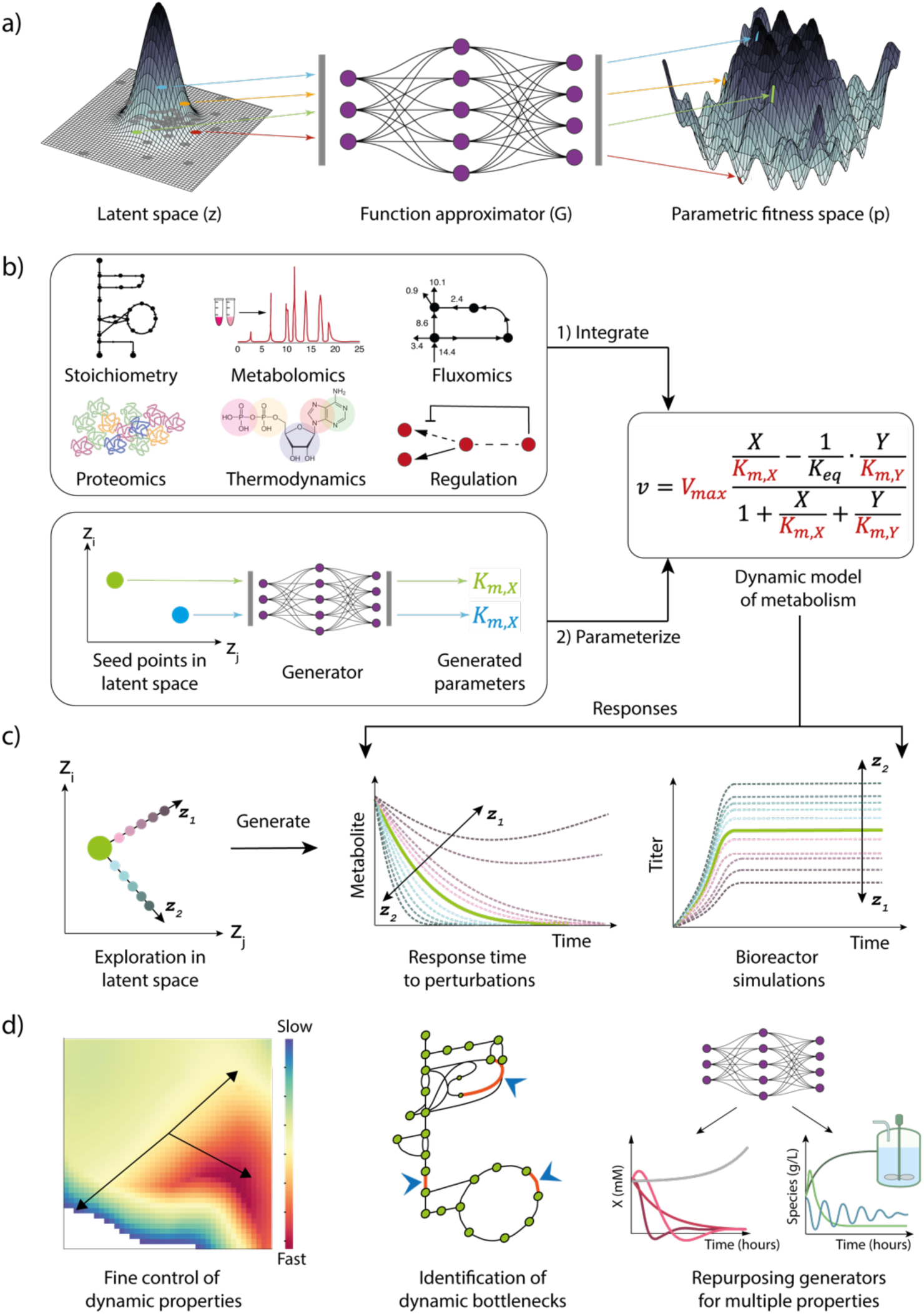
The framework for exploring the dynamic landscapes of metabolism through latent space. **(a)** Nonlinear function approximators, e.g., generative neural networks (G), map a complex parametric space (p) to a simpler latent space (z). **(b)** Generative learning-based methods produce kinetic parameters that satisfy desired dynamic properties, after relevant quantitative and structural information has been integrated. **(c)** Exploring around a latent seed point of a kinetic parameter set allows for generating novel parameter sets giving rise to dynamic models with distinct dynamic properties **(d)** A few applications of the methodology presented in this paper.

## RESULTS

### Navigating dynamic properties of metabolic networks

Metabolic networks are characterized by their complex topology and intricate dynamics, with metabolic reactions from different pathways operating across a range of time scales.^44,45^ Reactions in glycolysis, the pentose phosphate pathway, the citric acid cycle, and the electron transport chain operate with short time constants. Conversely, protein synthesis, DNA replication, and RNA transcription exhibit longer time scales, reflecting the time-intensive process of their molecular assembly. A fundamental requirement of dynamic models based on mechanistic rate laws is to capture such dynamic properties of metabolism while aligning with experimental observations.^14,19^ To meet this requirement, we explored the latent space of generator neural networks and investigated how to systematically create biologically relevant dynamic models — models that capture the time scales and behaviors of metabolic processes (Methods). We quantified the dynamics of the parameterized kinetic model using the dominant time constant, *τ_max_*. The dominant time constant corresponds to the slowest decaying mode in a system’s response, meaning this time constant has the greatest impact on the long-term evolution of the system dynamics (Methods).

To this end, we utilized the REKINDLE framework to train generators that produce biologically relevant dynamic models of *E. coli* central carbon metabolism, parameterized by 259 independent kinetic parameters (Methods, Supplementary Figures 3 and 4). After training, we selected a generator with a high incidence of biologically relevant models to generate 100 kinetic parameter sets, 80 of which were biologically relevant, using a 127-dimensional Gaussian vector as the latent input. We randomly chose one of the 80 models and explored how modifying its corresponding latent input affects the dynamic properties of the resulting parameterized metabolic network (Figure 1c). As a first step, we increased each latent feature by 50% individually and assessed its impact on the resulting generated kinetic parameter set. We found that even a single latent feature change altered multiple kinetic parameters due to the nonlinear behavior of the neural network (Figure 2a, Methods). Some latent features had a more pronounced effect on the model than others, whereas some did not impact the kinetic parameters at all. This suggests that only a subset of latent features plays a significant role in shaping kinetic parameters, while others may encode different aspects of the system, such as nonlinear behaviors or structural properties of the metabolic network.

Next, we examined how the dominant time constants, *τ_max_*, of the parameterized system change as we alter each latent feature. We found that altering each latent feature affected *τ_max_* (Figure 2b). Some changes made the model faster (*τ_max_* decreased), while others made it slower (*τ_max_* increased). We also tested different levels of latent feature modification (10%, 25%, and 50%) and observed the change in *τ_max_* with the modification magnitude (Supplementary Fig 5). Interestingly, the effect on *τ_max_* reversed when latent features were decreased by 50% compared to when they increased by 50% (Supplementary Fig 6). For example, increasing latent feature 22 by 50% decreased *τ_max_* by 14%, while decreasing it by 50% increased *τ_max_* by 13%.

**Figure 2.**
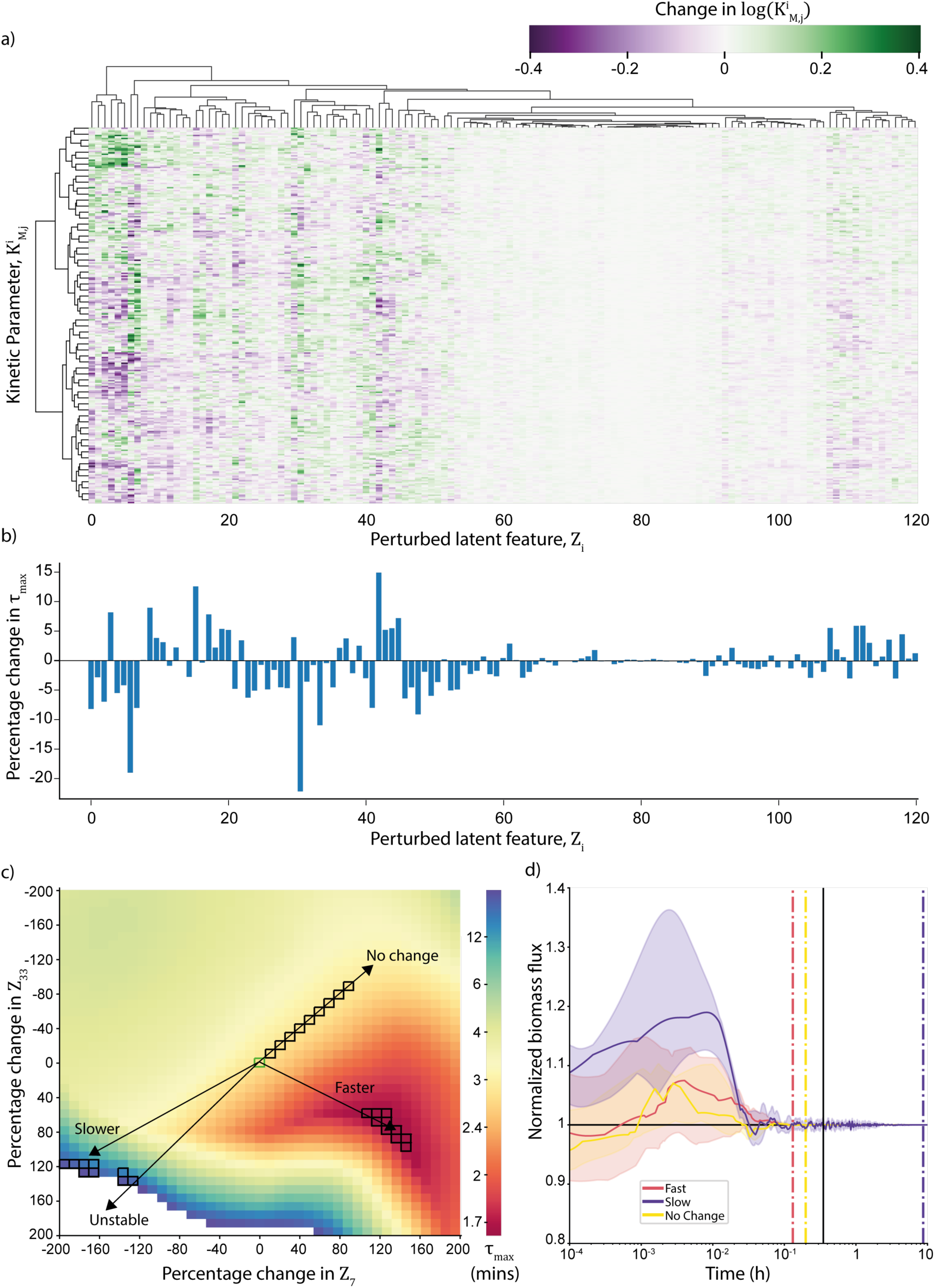
Latent space augmentation enables control of metabolic network dynamics. **(a)** Change in all log-transformed *K_M_* values from the original model when each of the 127 latent features is individually increased by 50% (Methods). **(b)** Corresponding percentage change in *τ_max_* from original model when all 127 latent features are increased by 50% one by one. **(c)** *τ_max_* values of the resulting models when the 2 most effective latent features that decreased *τ_max_* are modified between ±200% from the original model simultaneously. **(d)** Dynamic responses of normalized biomass flux of chosen models from Figure 2c (black squares). The black line indicates the doubling time of wild-type *E. coli*. Color-dashed lines represent time points when the median response returns within 5% of the reference steady-state value.

This reversal was consistent across all latent features, suggesting that modifying latent features in a neural network-generated kinetic model enables easy control of dynamic responses.

To further investigate this observation, we selected the two latent features with the highest impact on *τ_max_* (Figure 2b), which make the model dynamics faster. We perturbed these features simultaneously, varying them between ±200% in 5% increments, and analyzed *τ_max_* of the resulting 1681 models (Figure 2c). The results indicate that the dynamics of the models change based on different combinations of perturbations — some becoming slower, others faster, some exhibiting local instability, while some remaining unchanged — depending on the direction within the 2-dimensional latent space (Figure 2c). To validate this, we selected the 20 fastest, 20 slowest, and 20 models with no dynamic changes from the 1681 models (Figure 2c, squares). We perturbed the steady metabolite concentrations by up to ±50% and analyzed the response times of the selected models (Methods). The non-linear responses confirmed the conclusions obtained by analyzing the dominant times. The fastest models, achieved through latent space manipulation, settled back to the reference steady state of biomass flux much quicker (7.8 mins, Figure 2d, red dashed line) compared to the models with no dynamic changes (11.9 mins, yellow dashed line) and the slowest models (522.5 mins, purple dashed line). Similar trends were observed for ADP, ATP, and NADH concentrations (Supplementary Figure 7). This demonstrates that simple and systematic transformations in the latent space of a trained generator can reliably generate models with diverse dynamic properties and bias these properties towards specific requirements. A comparative discussion of latent space exploration and variance-based global sensitivity analysis^46,47^ is presented in Supplementary Note 3.

### Discovering dynamic bottlenecks in metabolic networks

Altering individual latent features provides a systematic and precise means to control the dynamic properties of a metabolic model. Yet, to gain a mechanistic understanding of metabolic processes, one must identify kinetic parameters that dictate the dynamic behavior of the metabolic network, particularly *τ_max_*, in any given parameterized model. To identify these kinetic parameters, we first examined how individual kinetic parameters responded to changes in the two most important latent features (Figure 2c) and found that all parameters varied smoothly in response to these modifications (Supplementary Figure 8). Next, we computed correlations between *τ_max_* and all 259 individual kinetic parameters, 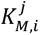, across the 1681 models from Figure 2c (Methods). Many kinetic parameters showed strong correlation or anti-correlation with *τ_max_* (Figure 3a, left), whereas such strong correlations were absent for models generated through random sampling in the latent space (Figure 3a, right). This difference arises because modifying key latent features with the highest impact on *τ_max_* alters the kinetic parameters that govern *τ_max_* in a systematic manner, whereas random sampling does not ensure proximity in parameter space, leading to arbitrary variations in *τ_max_* values.

**Figure 3.**
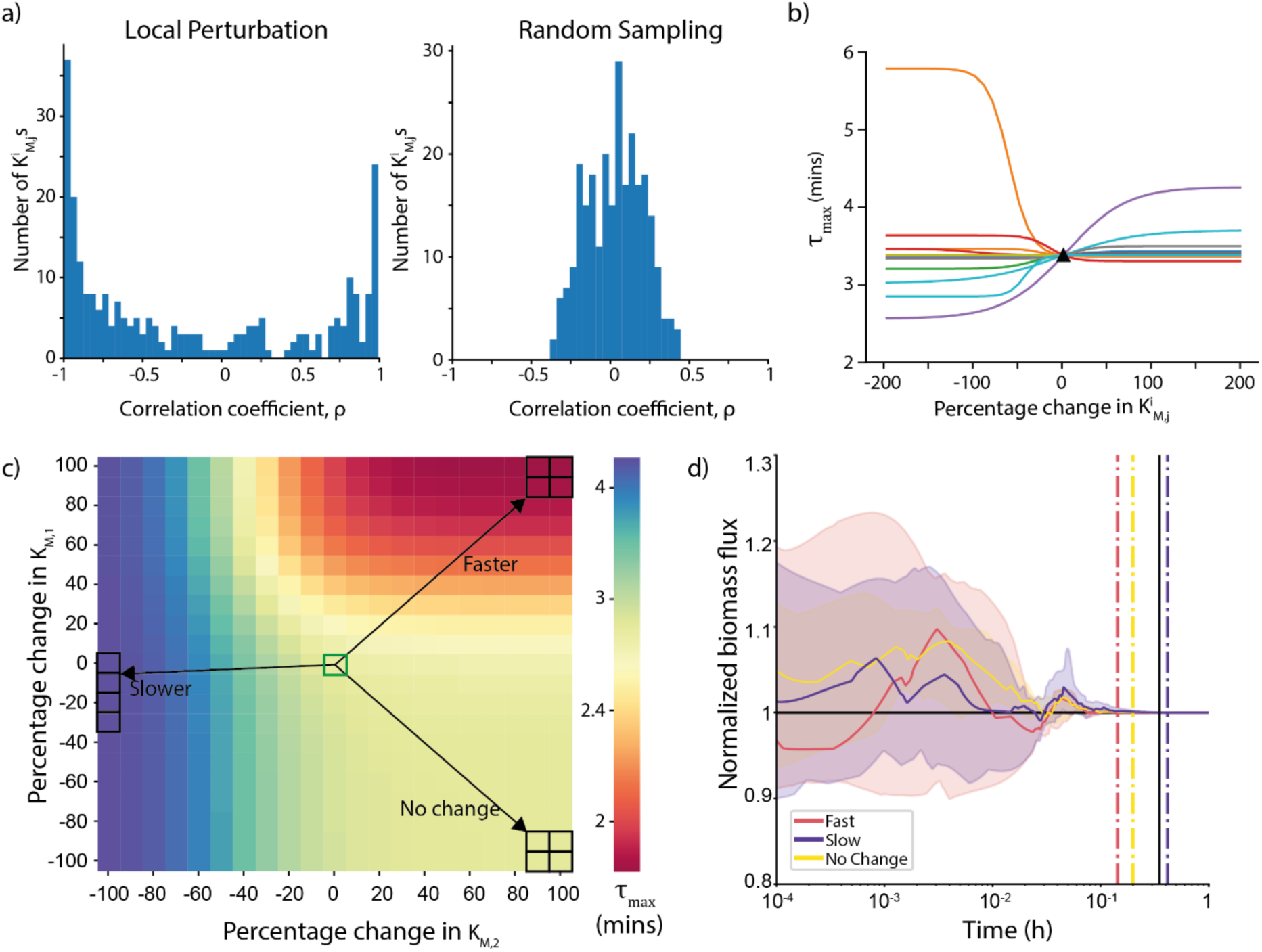
Enhancing interpretability at the parametric level with latent space augmented datasets. **(a)** Spearman Correlation coefficients between *τ_max_* of the parameterized system and all 259 kinetic parameters (left) of all 1681 models obtained in Figure 2c (right) when 100 models are randomly sampled from the generator **(b)** *τ_max_* values when 10 highest correlated/anti-correlated) parameters from 3a are discretely perturbed from the values of the original model. Black triangle represents *τ_max_* the original model. **(c)** *τ_max_* of resulting models when the top 2 parameters with highest impact on *τ_max_* in 3b are perturbed simultaneously from the original model. **(d)** Dynamic responses of normalized biomass flux of chosen models from Figure 2c (black squares). The black line indicates the doubling time of wild-type *E. coli*. Color-dashed lines represent time points when the median response returns within 5% of the reference steady-state value.

Next, we selected the 10 kinetic parameters with the strongest correlation or anti-correlation with *τ_max_*and individually varied each of them between ±100% in 10% increments to observe their impact on the *τ_max_* values. Interestingly, we observed that not all parameters that exhibit strong correlations, had a substantial effect on *τ_max_* when perturbed individually (Figure 3b). That is, some of these parameters are sloppy^48^, i.e., *τ_max_* is insensitive to changes in these parameters, and their apparent correlations with *τ_max_* do not reflect a causal influence. The observed correlations are instead a consequence of the nonlinear mapping learned by the generator neural network, which transforms points in the latent space into kinetic parameter sets. As a result, when a latent feature is modified, multiple kinetic parameters are adjusted proportionally, even if some of them do not individually influence *τ_max_*. This highlights the importance of decoupling correlation from causal influence, especially in complex, nonlinear generative models. The performed analysis allows us to distinguish between parameters that are merely correlated due to the latent space structure and those that truly govern the model’s response speed (Figure 3b).

Based on this analysis, we identified the two parameters with the highest impact on *τ_max_* by measuring the total deviation from the original model’s *τ_max_* (Figure 3b, black triangle), filtering out sloppy parameters that vary widely without affecting *τ_max_*. We then perturbed these two parameters, 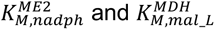, simultaneously between ±100% in 10% increments and examined the resulting *τ_max_* values (Figure 3c). Similar to the results in Figure 2c, we found that modifying only these 2 out of 259 parameters significantly changed *τ_max_*, thereby affecting the overall dynamic response time of the entire metabolic network.

To validate this finding, we again performed nonlinear integration of the 20 fastest, 20 slowest, and 20 models with no dynamic changes by perturbing the reference metabolite concentrations by up to ±50% and analyzing their response times (Figure 3c, squares). The fastest models returned to the reference steady state of biomass flux much sooner (8.6 mins, Figure 3d, red dashed) compared to the models whose dynamics did not change (11.9 mins, yellow dashed), and the slowest models (24.9 mins, purple dashed). Similar trends were observed for ADP, ATP, and NADH concentrations (Supplementary Figure 9), demonstrating that latent space manipulation enables direct identification of the key parameters governing a model’s dynamic properties. In this case, 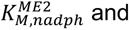 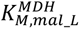 primarily control 𝜏*_max_* of the parameterized dynamic model.

Since kinetic parameters are inferred to match experimentally measured steady-state metabolite concentrations and fluxes, one can yield different sets of kinetic parameters for different reference steady states. Consequently, 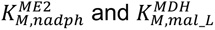 do not necessarily control 𝜏*_max_* in every model, as the dynamic properties of metabolic networks arise from complex interdependencies among kinetic parameters. Thus, for each set of kinetic parameters and the corresponding reference state of concentrations and fluxes, different parameters may emerge as key controllers of *τ_max_*. To address this, we next investigated whether certain parameters or enzymes consistently act as dynamic bottlenecks in *E. coli* central carbon metabolism both within models satisfying a given steady state and across models from multiple steady states.

To this end, we sampled four different thermodynamically feasible reference profiles of fluxes and metabolite concentrations using Thermodynamic Flux Balance Analysis^49^ (Methods). For each reference state, we employed REKINDLE to obtain generators providing biologically relevant dynamics. We then generated 100 models per state and selected relevant models with *τ_max_* ≤ 6.67 mins. Finally, for each relevant model, we repeated the analysis from Figures 2 and 3, where we: (i) perturbed latent space features individually, (ii) identified those with the highest impact on *τ_max_*, and (iii) determined the top 3 kinetic parameters using correlation and filtering from the resulting set of models. We focused on the top 3 parameters, as those beyond them had a negligible impact on *τ_max_* in most cases (Figure 3b). We then identified enzymes whose associated kinetic parameters consistently affected *τ_max_* across all steady states, with key reactions in the TCA cycle and glycolysis playing a dominant role (Supplementary Figure 10). Indeed, kinetic parameters associated with Pyruvate Dehydrogenase (PDH), Glyceraldehyde-3-Phosphate Dehydrogenase (GAPDH), and Citrate Synthase (CS) primarily determined *τ_max_* across the network (Figure 4a).

**Figure 4.**
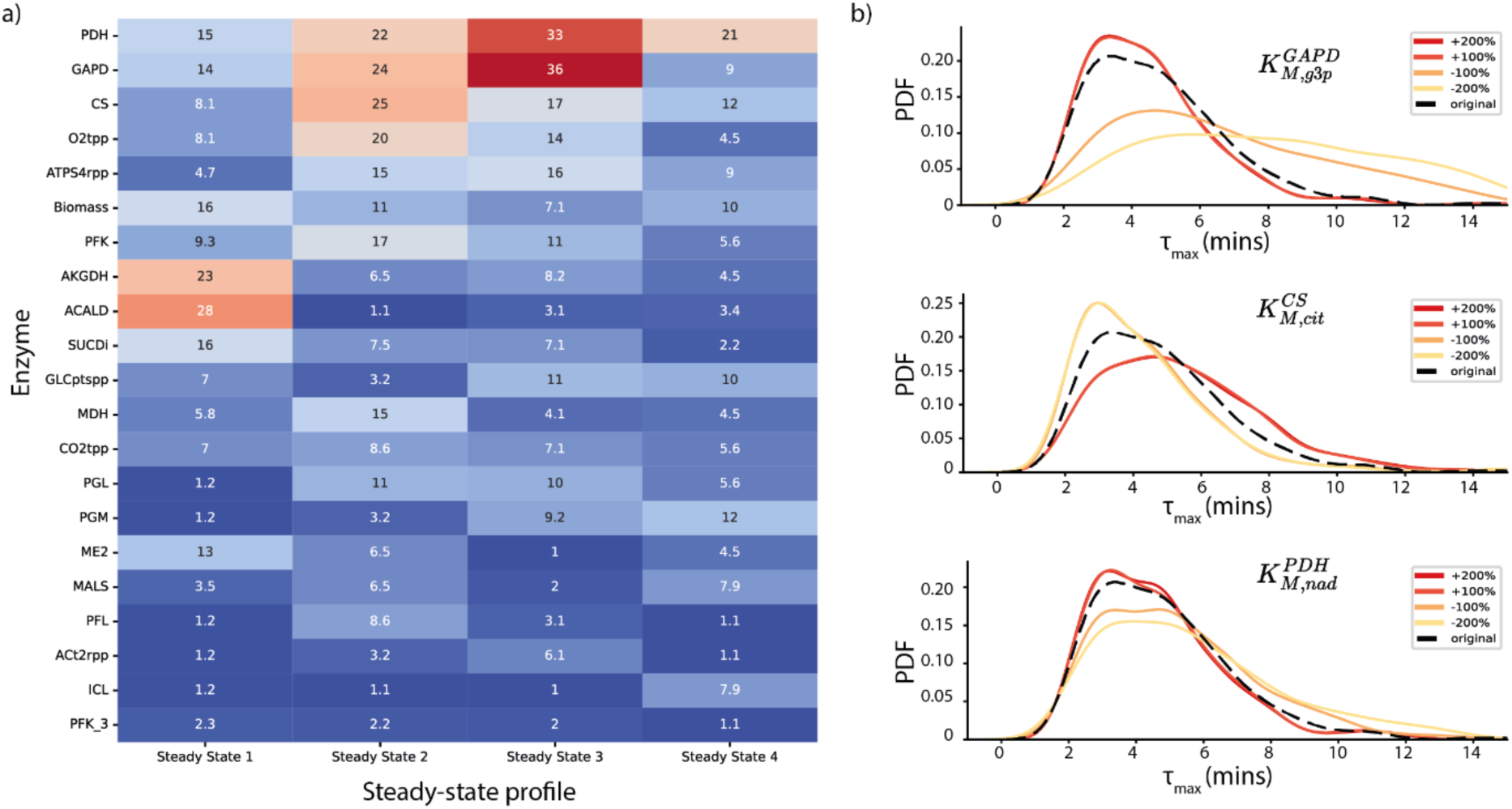
Discovering general dynamic bottlenecks in the central carbon metabolism of *E. coli*. **(a)** Occurrence of enzymes (in %) whose associated kinetic parameters significantly affect *τ_max_* of the E. coli central carbon metabolism across 4 different steady-state profiles. Red color denotes high occurrence, while blue indicates low occurrence. **(b)** Distribution of *τ_max_* across models when 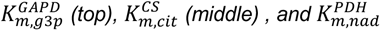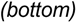 are perturbed from their original values by different magnitudes (colored lines). The black dashed line represents the *τ_max_* distribution of the original, unperturbed models.

To substantiate this result, we modified the kinetic parameters of these 3 enzymatic reactions one at a time and evaluated the *τ_max_*of the resulting models. We found that changing specific parameters, such as 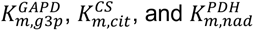, consistently affected the 𝜏*_max_* of all models across all steady states. In contrast, alterations to other parameters within the same metabolic reactions had little to no effect (Supplementary Figure 11). Taken together, these results indicate that 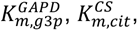 and 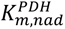, consistently emerge as dynamic bottlenecks governing 𝜏*_max_* across the ensemble of models and steady states we examined. Because the parameter ensembles were generated with REKINDLE, which recreates the same parameter distributions obtained through unbiased sampling while enforcing consistency with experimental data, this pattern is unlikely to be an artifact of the generator itself. We also examined whether such important enzymes co-occurred for any particular steady-state profile, but we did not find strong evidence of this (Supplementary Note 2).

### Robustness and generalizability across organisms and generative models

To assess the generalizability of our findings beyond *E. coli* and to evaluate robustness across generators exhibiting different incidences of desired dynamic properties, we performed an additional analysis using multiple generative models trained to parameterize large-scale kinetic models of *S. cerevisiae*. Specifically, we employed the RENAISSANCE framework to train generators capable of producing biologically relevant dynamic models of *S. cerevisiae* central carbon and aromatic amino acid metabolism, parameterized by 1,098 Michaelis constants and 303 maximal reaction velocities (Methods).

From the generators obtained at different stages of the training process, we selected three corresponding to early, intermediate, and late stages of training, characterized by increasing incidences of biologically relevant outputs (5%, 15%, and 23%, respectively; Figure 5a). For each of the three generators, we sampled 500 kinetic parameter sets. From each resulting population, we selected three representative kinetic models closest to the 10th, 50th, and 90th percentiles of the dominant eigenvalue distribution (Figure 5c) and examined how perturbations of their latent inputs affect model dynamics. This design yielded nine cases spanning generators of varying quality and distinct regions of latent space explored around representative latent input points (Figure 5b,c).

Across these cases, the two latent features most strongly influencing the dominant timescale *τ_max_*depended on both the nonlinear generator mapping and the selected latent input point, and varied substantially across training stages (Figure 5b, Supplementary Figure 19). Similarly, the structure of the latent space differed across generators, with no simple or universal relationship between *τ_max_* and either generator quality or the location of the latent input point. Despite this heterogeneity, latent space exploration consistently enabled both acceleration and deceleration of model dynamics, corresponding to decreases or increases in *τ_max_*.

For example, for the late-stage generator and a latent input point near the 90th percentile (Figure 5b, bottom right), the original model exhibited *τ_max_* ≈ 13ℎ. Perturbation of the two most effective latent features within ±200% allowed the dynamics to be accelerated to approximately 1.71ℎ (∼7.6-fold decrease) or slowed to beyond 25ℎ (at least ∼1.9-fold increase). In contrast, for the early-stage generator and a latent input point near the 10th percentile (Figure 5b, top left), the accessible dynamic range was more limited: *τ_max_* could be reduced from approximately 2.75ℎ to 2ℎ (∼1.4-fold decrease) or increased to about 3.7ℎ (∼1.3-fold increase) under equivalent perturbations. In addition, for some configurations, particularly for the intermediate-stage generator, specific combinations of latent input values led to locally unstable dynamics, manifested as regions of instability in latent space (Figure 5b, white regions). Importantly, this latent-space-based modulation enables systematic control of dynamic behavior without the need to individually modify or re-fit each kinetic parameter in the model, which would be computationally intractable.

**Figure 5:**
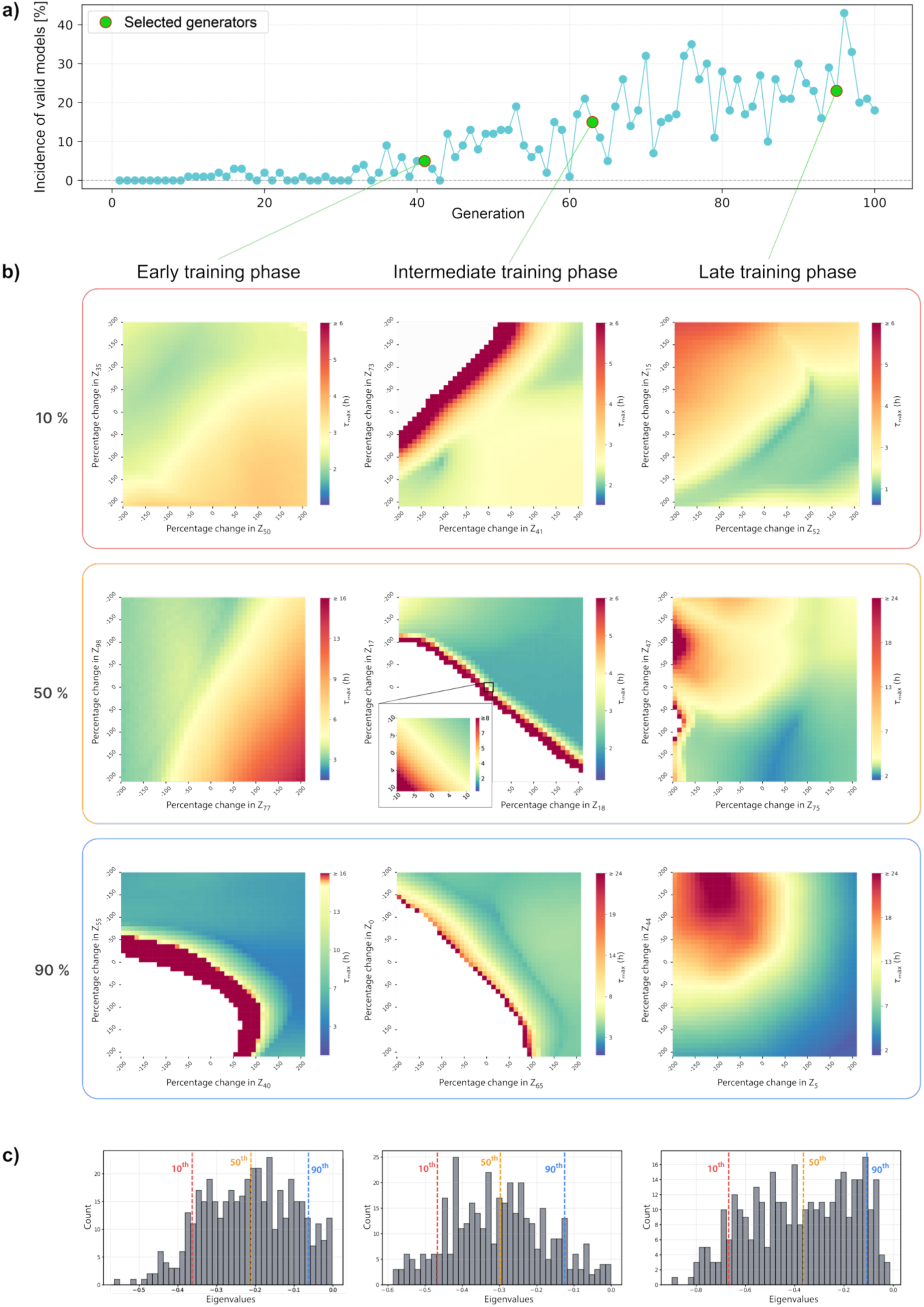
Latent Space Control of Metabolic Network Dynamics in *S. cerevisiae*. **(a)** The fraction of models exhibiting desired dynamic properties increases over training. Generators from early, intermediate, and late training phases were selected for subsequent latent space exploration (green circles). **(b)** Heatmaps of *τ_max_* values for nine representative cases obtained by perturbing the 2 most effective (Top 2) latent features within ±200% of their original values. Along the horizontal axis, generators from different training phases were used, as defined in (a). Along the vertical axis, latent points closest to the 10^th^ (red), 50^th^ (yellow), and 90^th^ (blue) percentiles of the model distribution shown in **(c)** were selected. The inset (center middle) shows a magnified portion of the latent space within ±10% of the original latent values. White regions indicate models with locally unstable dynamics. Each of the nine cases corresponds to a distinct pair of Top 2 latent features. **(c)** Distribution of dominant eigenvalues for models generated by the three selected generators, with the 10^th^ (red), 50^th^ (yellow), and 90^th^ (blue) percentiles indicated.

Taken together, these results demonstrate that latent space exploration provides a consistent mechanism for tuning the dynamic properties of large-scale kinetic models, even when starting from generators trained at early stages with relatively low incidences of biologically relevant outputs. The extent of achievable modulation depends on both generator quality and local latent space structure. We note, however, that the range of attainable dynamic behaviors is ultimately bounded by the expressivity of the trained generator and the structure of its learned latent manifold. Notably, generators obtained at later training stages exhibit smoother latent landscapes and substantially greater flexibility in the range of attainable dynamic behaviors, indicating the importance of adequate generator training for effective latent control of metabolic network dynamics.

### Repurposing generators for multiple dynamic properties

Dynamic models of metabolism often need to satisfy multiple dynamic requirements to facilitate applications in systems and synthetic biology design and testing. One common application of dynamic models is in predicting cellular responses in bioreactors^50^, where cells are used as ‘biofactories’ to produce a non-native compound or overproduce a native compound. Kinetic models need to accurately capture experimental fermentation curves in terms of growth, yield, and titers before being used for optimizing strain design. A recent study^14^ showed that only a small fraction of models obtained by traditional sampling-based kinetic modeling frameworks (212 out of 91,852 models, i.e., 0.23%) successfully capture dynamic properties of metabolism and experimental fermentation data. Moreover, this fraction decreased even further when sampled models were tested on different recombinant strains in addition to wild-type physiology. Therefore, deriving models that simultaneously satisfy multiple dynamic properties or multiple physiologies requires considerable computational resources and systematic downstream processing. Here, we aimed to leverage latent space manipulation to repurpose trained generators – originally designed for a single dynamic property – to construct models that satisfy multiple dynamic requirements.

**Figure 6:**
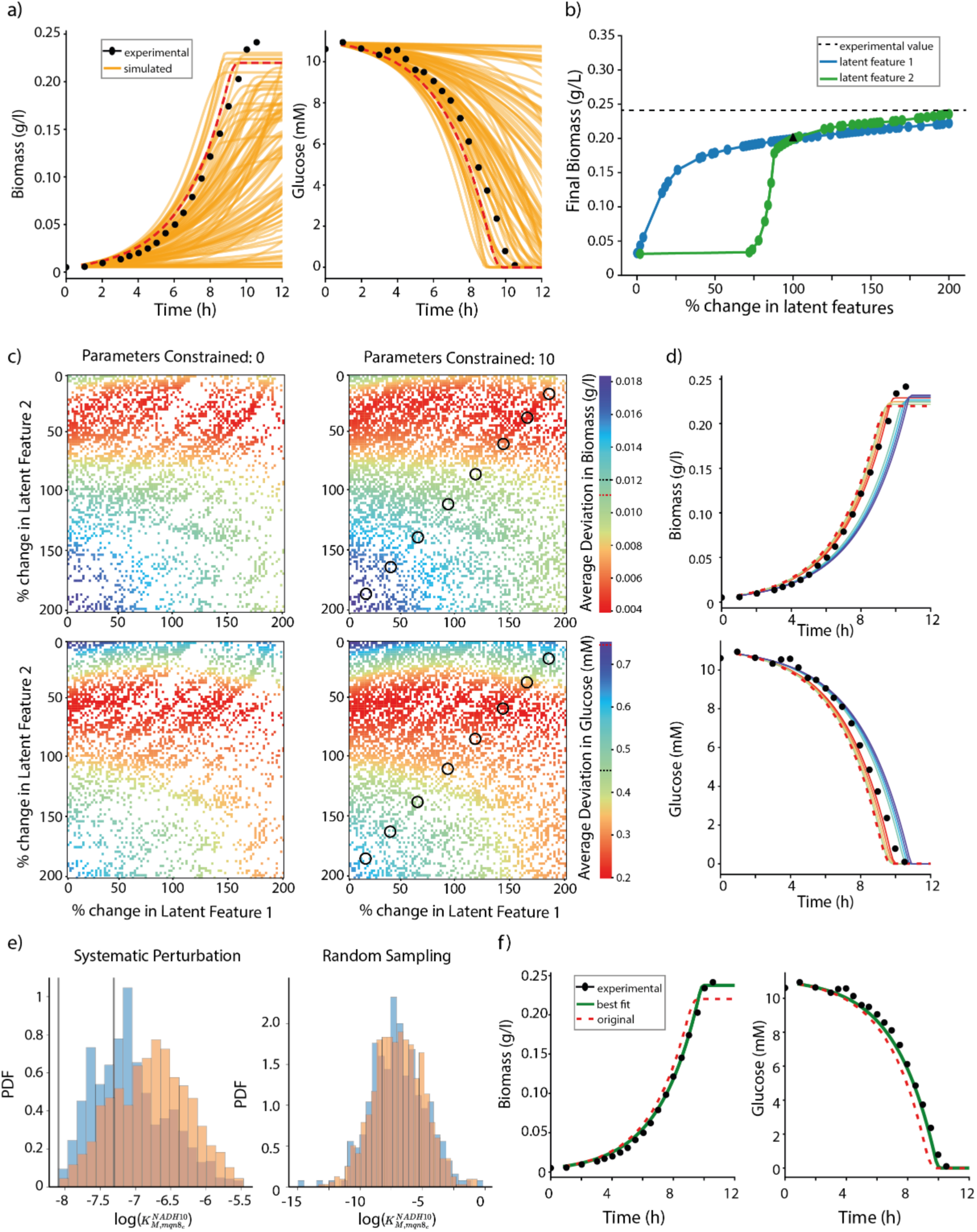
Repurposing generators to enforce multiple dynamic properties of metabolic models. **(a)** Bioreactor responses of 300 randomly sampled models generated by a neural network trained to capture the relevant linearized dynamics of anaerobic *E. coli* metabolism, for Growth/Biomass (left) and Glucose (right). The response marked in dashed red is chosen for further studies **(b)** Final biomass titers of model responses when two most important latent features are perturbed from their original values. The black triangle represents the final biomass titer of the original model (red line from Figure 6a) **(c)** (left) Average deviations of simulated responses of biomass (top) and glucose (bottom) from experimental bioreactor datapoints when the two latent features in Figure 6b are perturbed simultaneously. (right) Average deviations of simulated responses of biomass (top) and glucose (bottom) from experimental bioreactor data points when the two latent features in Figure 6b are perturbed simultaneously, with constraints imposed on 10 stability-relevant parameters (see main text and Figure 6e). The dashed black line on the color bar represents average deviations of predicted responses for the same system as obtained by Varma et al^51^ **(d)** Bioreactor responses of 8 models sampled from Figure 6c, Right panel, black circles. The responses are colored according to the corresponding sampling space as mapped in the color bar in Figure 6c. The red dashed line represents the original model response. **(e)** Distribution of the parameter 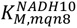 in generated stable (blue) and unstable (orange) models. The models were obtained either by systematic perturbation of the latent point of the model in Figure 6a (left) or by random sampling of the latent space of the trained generator (right). Gray vertical lines in the top panel represent the 30^th^ percentile values for this parameter that favors stability. NADH10: NADH dehydrogenase (Menaquinone-8, 0 protons), mqn8_c: Menaquinone-8. **(f)** The bioreactor response of the best model (green) obtained through latent space exploration, and the original model response (red dashed).

Building on this approach, we demonstrate how generators trained to produce models with valid *τ_max_*can be repurposed to generate models that satisfy specific fermentation data for anaerobic *E. coli* using latent space manipulation. We used RENAISSANCE^19^ to train generator neural networks to generate models with valid linearized dynamics for the anaerobic *E. coli* model (Methods, Supplementary Figure 12). Selecting the generator with the highest incidence of valid models (30%), we generated 1000 models. After integrating initial experimental conditions, we performed bioreactor simulations on the 300 valid models (Methods). While we observed a range of responses, none of the 300 models perfectly fit the experimental data (Figure 6a). This was expected, as the generator was not explicitly trained to produce models satisfying this specific fermentation dataset.

To obtain models that more closely fit the experimental data, we selected one that approximated the observed trends but exhibited lower final biomass titers and a faster glucose uptake rate (red dashed line, Figure 6a). Using its corresponding latent space input, we investigated whether latent space manipulation could generate models with improved agreement. To quantify the goodness of fit, we calculated the average deviation (E) between simulated responses and experimental data, following Varma et. al^51^ (Methods). The average deviation of the chosen model (*E\**) wa 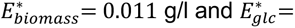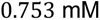). As a first step in exploring latent space, we modified each of the 99 latent features of the chosen model one at a time. Each feature was perturbed between ±100% in 1% increments, resulting in 101 modified models per feature. We then simulated the bioreactor responses for all modified models and analyzed the final biomass titers. Consistent with previous findings, perturbing certain latent features significantly affected the final biomass titers, while most had little to no impact (Supplementary Figure 13). However, unlike in previous studies, we observed that some models became locally unstable upon latent feature perturbations.

Given that fitting the biomass titers was more challenging than fitting the glucose uptake rate, we selected the two latent features with the highest impact on the biomass titers (Figure 6b). Since increasing these two features also increased final biomass titers, we perturbed them between 0 and 200% of their original values in 1% increments (Figure 6b). Among the generated 10201 new models, 29% had valid *τ_max_* (Methods). We then performed bioreactor simulations for the valid models and evaluated their average deviations from experimental values (Figure 6c, left). Although 71% of the models were unstable, we observed distinct trends in goodness-of-fit depending on the perturbation levels in the selected latent features.

Next, we investigated whether the instability of perturbed models could be reduced by comparing the distributions of kinetic parameters from two sources: (i) random sampling of the generator’s latent space (1,000 models, Figure 6a), and (ii) systematic perturbation of latent space inputs (10,201 models, Figure 6c). We found that parameters derived from systematic perturbation provided better delineation between stable and unstable models than those obtained through random sampling (Supplementary Figure 14). An example of this delineation for 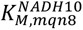 is shown in Figure 6e. To quantify these differences, we ranked all 478 kinetic parameters by measuring the divergence between stable and unstable model distributions using Jensen-Shannon (JS) entropy (Methods). The 10 parameters with the highest distribution differences were constrained within ranges that favor stability in the 10201 perturbed models (Figure 6e, vertical gray lines). Constraining these 10 parameters increased the local stability incidence from 29% to 47% and improved the resolution of the original perturbed set of models (Figure 6c).

We then repeated this procedure by constraining the top 20, 50, 100, 150, and 200 parameters, and observed that local stability increased up to 60% (Supplementary Figure 15). With improved stability through latent space analysis, we next examined the bioreactor responses of the constrained models (Figure 6c, right; Supplementary Figure 16, 17). Many constrained models exhibited a better fit than the original unperturbed model (Supplementary Figure 16, 17). Indeed, constraining just 10 parameters not only yielded better fitting models but also introduced a range of responses with different dynamics and final biomass titers, enabling finer tuning (Figure 6d). Increasing the number of constrained parameters further improved model performance, with the best-fitting model obtained by constraining 200 parameters (Figure 6f, green), achieving an average deviation *E^best^*, where 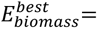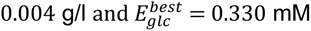, representing a 63% and 56% improvement in goodness-of-fit for biomass and glucose predictions, respectively, compared to the original model. These findings demonstrate that the latent space of neural networks can be leveraged to repurpose generators trained for a specific dynamic property, enabling them to satisfy additional constraints without requiring explicit retraining.

## DISCUSSION

Kinetic models of metabolism allow us to integrate and consolidate data from cellular processes and predict transient and steady-state metabolic responses in a quantitative and deterministic manner. This capability makes them invaluable tools for numerous applications in biomedical and biotechnological sciences. Recently developed data-driven generative methods, such as REKINDLE and RENAISSANCE, have significantly improved the efficiency of kinetic model parameterization, enabling dynamic studies across multiple physiologies and large patient cohorts. These methods rely on predefined network stoichiometry, kinetic mechanisms, and feasible steady states, but do not require exhaustive kinetic measurements for individual parameters. As the applications of these methods will inevitably extend to other domains of biotechnology and health, research efforts in this field would greatly benefit from the ability to repurpose generative neural networks designed for a specific physiology and/or organism to other studies. This would allow researchers to study new physiologies and organisms without having to undergo the lengthy modeling process again. The methodology proposed in this paper aims to address this challenge by using the latent space of neural networks employed in these advanced methods to repurpose generators, allowing them to capture diverse dynamic properties while enhancing interpretability.

As computational and modeling techniques continue to advance, and as increasingly rich datasets from omics and other high-resolution experimental methods become available, *in silico* models will increasingly align with real biological systems. These models will provide fundamental insights while reducing reliance on extensive experimental studies. Generative neural networks inherently encode higher-level or emergent system properties in their latent space during training. By manipulating these latent embeddings, we can systematically access and control these properties. Moreover, latent space encodings facilitate model reduction by decreasing the dimensionality of the parametric space, enabling fine-tuned control over emergent dynamic properties using simple linear transformations while maintaining reference steady-state conditions. The effectiveness of this representation depends primarily on the consistency of the network definition and the availability of feasible steady states, rather than on comprehensive kinetic parameter datasets. These transformations generate structured datasets that are well-suited for statistical analysis, allowing researchers to identify key system parameters or parameter combinations that govern network-level properties—an approach not feasible with conventional random parameter sampling in kinetic model parameterization.

We demonstrate how the latent space exploration of generative neural networks can offer system-level insights valuable to both modelers and experimentalists alike. For example, simple latent space transformations can help identify candidate dynamic bottlenecks in metabolic networks (Figure 4), suggesting points for targeted experimental interventions without requiring complex downstream analyses. Importantly, these insights reflect the region of parameter space consistent with the observed experimental data and may evolve as broader physiological conditions are considered. Furthermore, repurposed generators that produce models satisfying multiple dynamic criteria can serve as robust starting points for *in silico* and experimental hypothesis testing, thereby extending the utility of these methods beyond their standard “off-the-shelf” application.

Importantly, the limits of applicability of latent space exploration are not determined by species identity per se, but by the consistency of the underlying metabolic model and the availability of feasible steady states used during generator training. For example, in the *S. cerevisiae* case, whose metabolism is comparatively less characterized at the kinetic level than that of *E. coli,* reproducible control over dominant time scales and stability properties is retained across generators and training stages, despite increased uncertainty in kinetic parameters. Differences in physiological conditions or organisms are reflected as distinct regions of latent space, which can be systematically explored and repurposed without retraining the generator, provided that the network structure and operating regime remain compatible with the training distribution. By contrast, arbitrarily large extrapolations, such as substantially altered network topology or incompatible steady-state regimes, are not expected to be captured reliably by a generator trained on a fundamentally different system. In this sense, the practical limits of applicability are set by the coverage of the training distribution and the structural assumptions encoded in the metabolic model, rather than by the latent space methodology itself.

We further demonstrate that latent space exploration can repurpose generators trained for a single property to produce models satisfying multiple dynamic constraints. This repurposing can, in principle, be extended to increasingly complex properties, thereby significantly reducing the computational costs associated with pruning large model ensembles for specific traits. Such an approach could streamline *in silico* strain design pipelines like NOMAD^14,15^. Because the approach exploits system-level structure learned during training rather than relying on precise identification of individual parameters, it remains applicable to non-model organisms and emerging genome-scale reconstructions with limited kinetic data. Moreover, since this methodology is model-agnostic, it can be applied to any dynamical system for which trained generators are available. At a minimum, a meaningful application requires a curated stoichiometric network, plausible kinetic mechanisms, and at least one feasible steady-state operating point. As generative machine learning methods like REKINDLE and RENAISSANCE become more widely used across metabolic studies and organism-specific designs, the growing availability of trained generators will further enhance the utility and accessibility of our proposed methodology.

## METHODS

### Dynamic models of *E. coli* and *S. cerevisiae* metabolism

#### *E. coli* central carbon metabolism

The kinetic nonlinear models employed in this study represent the central carbon metabolism of *Escherichia coli* in its wild-type form. These models are based on the models by Varma and Palsson^52^ and have been extensively analyzed using the SKimPy^53^ toolbox and the REKINDLE^31^ method. The model encompasses 64 metabolites, distributed between the cytosol and the extracellular space, and includes 65 reactions, of which 16 are transport reactions. Among the 64 metabolites, 15 are designated as boundary metabolites, as they are localized in the extracellular space. The remaining 49 metabolites are confined to the cytosol, where their dynamics are governed by a system of ordinary differential equations (ODEs) describing mass balances. Each reaction was assigned a kinetic mechanism based on its stoichiometry, and the system was subsequently parameterized using 411 kinetic parameters. Out of these 259 were concentration associated Michaelis Menten constants, K_M_s, and the remaining 152 parameters are maximal reaction velocity rates, V_max_s.

From this context-specific model, 4 steady-state profiles were sampled using thermodynamics-based flux balance analysis^49^ implemented with the pyTFA tool^54^. Each steady-state profile consists of metabolite concentrations, metabolic fluxes and thermodynamic variables. Once these profiles are sampled, we generate kinetic models for each steady state using RENAISSANCE or REKINDLE, as specified in the studies.

### Anaerobic metabolism of *E. coli*

The kinetic model was constructed using a reduced stoichiometric network of the central carbon metabolism of *E. coli* grown under anaerobic conditions. The reduced stoichiometry was built using redGEM^55^ and lumpGEM^56^ and includes core pathways like glycolysis, PPP, TCA, pyruvate metabolism and a lumped reaction for growth. When building the stoichiometry, we blocked the flux through oxygen uptake and any electron transport chain reaction that used oxygen as an electron acceptor to ensure the model accurately represented anaerobic conditions. The resulting steady-state model had 180 reactions, including 40 boundary reactions and 34 transport reactions. It included 141 metabolites in total, of which 104 were intracellular, and 37 were extracellular, corresponding to 119 unique metabolites.

The model was tailored to a specific context by imposing physiological and experimental data from the study conducted by Varma and Palsson.^51^ The growth rate was set to 0.4322 h^-1^ and the glucose uptake rate to 18.50 mmol gDW^-1^ h^-1^ (gram per weight per hour). Extracellular metabolites not reported as secreted or found in the media were constrained to their minimum feasible secretion/uptake rate, and a maximum possible concentration of 10^-4^ M, an order of magnitude lower than the reported concentrations of fermentation products. Known extracellular concentrations for media components were imposed with a +-20% error. For the primary fermentation products (ethanol, formate, acetate) the steady-state concentrations were also constrained to 10^-4^ M. Next, constraints were imposed on thermodynamic variables, calculated using the group contribution method^57,58^ to ensure that the sampled flux directionalities and metabolite concentrations were consistent with the second law of thermodynamics.

A total of 10,000 steady-state profiles, consistent with the integrated data, were sampled from this context-specific model using thermodynamics-based flux balance analysis implemented in the pyTFA tool.^54^ Each steady-state profile includes metabolite concentrations, metabolic fluxes, and thermodynamic variables. Once these profiles were obtained, we generated kinetic models around each steady state. We used the profile with index 3709 as the reference in the study “Repurposing generators for multiple dynamic properties” and applied RENAISSANCE for the parameterization.

The kinetic model was built with SkIMpy^53^ and comprises 63 mass balances and 138 rate laws with various mechanisms related to the associated reaction. Specifically, there are 21 Convenience mechanisms^59^, 55 Generalized Reversible Hill^60^, 2 Bi-Uni Reversible Hill^60^, 41 Reversible Michaelis-Menten, 18 Uni-Bi Reversible Hill^60^, and one multi-substrate Monod (Irreversible Michaelis-Menten) for the growth rate. These mechanisms are parametrized by 463 Michaelis constants, K_M_s, and 138 maximal velocities, V_max_s, totaling 601 kinetic parameters.

### S. *cerevisiae* core metabolism

To demonstrate the generalizability of the proposed framework across biological systems, we applied it to a more complex eukaryotic organism. Specifically, we considered a strain of S. *cerevisiae* (ST10284), which enabled direct reuse of a curated context-specific model and dataset from the study by Narayanan, Jiang, and Wang et al.^15^ The dataset contains 5,000 steady-state profiles for ST10284, generated using the pyTFA tool. Consistent with the procedure described by Narayanan, Jiang, and Wang et al., we selected the steady-state profile closest to the mean of all sampled profiles as the reference operating point for constructing the kinetic model. This profile corresponded to index 3191 in the sampled ensemble.

Physiologically plausible ranges for Michaelis-Menten constants (K_M_s) for S. *cerevisiae* were obtained from the BRENDA enzyme database, following the curation reported by Chang et al.^61^ These ranges were subsequently used as constraints during kinetic parameterization to ensure consistency with experimentally reported yeast enzyme kinetics.

### Valid dynamic responses of *E. coli* metabolism

Once the structure of the ODEs has been defined and steady state data has been integrated as outlined in the sections above, we need to parameterize the model to fully define it. Once parameterized, the model exhibits different dynamic behaviours depending on the exact values of the kinetic parameters, not all of which are biologically valid or physiologically desirable.

In this study, we define kinetic models as stable if they exhibit local stability in the reference steady state profile of fluxes and metabolite concentrations and if all characteristic times of the aperiodic model response remain within physiologically and biologically plausible limits. To assess local stability and time constants, we compute the Jacobian matrix of the dynamic system^23^. The stability of a given model is determined by the eigenvalues of the Jacobian: a model is considered locally stable if all eigenvalues have negative real parts, whereas the presence of any positive real part indicates instability.

For models classified as locally stable, we define the characteristic time constant, *τ_max_*, of the linearized system as the inverse of the real part of the largest eigenvalue of the Jacobian, λ_max_. These time constants provide insights into the model’s dynamic behavior. Fast metabolic processes, such as glycolysis and the electron transport chain, are associated with small time constants, whereas biosynthetic processes operate on longer timescales. The characteristic timescales must remain within biologically and physically relevant bounds.

To ensure physiological validity, we impose the constraint that the aperiodic model response should not exceed the timescale of cell division. Specifically, we enforce that all characteristic response times be at least three times faster than the cell’s doubling time, ensuring that metabolic perturbations settle within 5% of the steady-state level before the next division. Additional constraints on response dynamics are also considered, such as ensuring that biochemical responses occur on a timescale slower than proton diffusion within the cell. Models satisfying these criteria are expected to accurately capture the metabolic dynamics observed in an organism.

In this study, we classify models as valid or invalid based on these physiological constraints. For aerobic *E. coli*, the dynamic responses of our models should be at least three times faster than the wild-type doubling time (∼21 min)^62^; that is, the dominant time constant of the model’s responses should be smaller than ∼7 minutes. Similarly, anaerobically grown *E. coli* has a doubling time of ∼120 minutes^51^; thus the dominant time constant of the model’s responses should be under ∼40 minutes to be deemed valid.

### Training generators to produce dynamically valid models

In this study, we use deep generative learning frameworks like REKINDLE^31^ and RENAISSANCE^19^ to parameterize the ODEs describing the dynamic model. These methods require a fully defined mathematical structure of the metabolic system and the integration of context-specific multi-omics reference data prior to parameterization (Figure 1b). Once these structural and data constraints are established, the frameworks employ distinct optimization strategies: REKINDLE utilizes adversarial training^36^, while RENAISSANCE relies on Natural Evolution Strategies (NES)^63,64^. These methods train neural networks to generate kinetic parameter sets that ensure the parameterized model exhibits desirable dynamic properties while satisfying the reference data, as described in the section above.

REKINDLE was used to train generators for obtaining valid models of *E. coli* central carbon metabolism for all the studies showcased in Figure 2,3 and 4.

### Training data for REKINDLE

REKINDLE requires labelled kinetic parameter sets from traditional sampling-based frameworks as training data. For this purpose, 5000 parameter sets were sampled for each of the 4 steady state profiles using the ORACLE framework^24,65,66^ . The training set had an incidence of 48%, 15%, 28% and 22% valid models for the 4 steady states, respectively. This training data was log-transformed and labelled as valid or invalid as described in the previous section before training.

### REKINDLE training

REKINDLE was employed on the training data to generate parameter sets with valid dynamic responses separately for each steady state. 3 statistical repeats were done for each steady state with randomly initialized generator and discriminator. For every repeat, training was deemed complete and successful when the generator and discriminator losses had stabilized and the discriminator accuracy reached ∼50% (Supplementary Figure 1), indicating the discriminator was not able to distinguish between samples from the training data and the generator’s samples. The incidence of valid models throughout the training process was recorded (Supplementary Figure 2) and the generators with highest incidences across the repeats were chosen for conducting further studies for each steady state.

RENAISSANCE was used to train generators for obtaining valid models of anaerobically grown *E. coli* metabolism for the studies showcased in Figure 6.

### RENAISSANCE training

RENAISSANCE was employed with following hyperparameters: generator population, 𝑛 = 20, search radius, 𝜎 = 10^−2^, learning rate, 𝛼 = 10^−3^, and learning rate decay, 𝑑 = 5%. The optimization was done with 10 statistical repeats with a randomly initialized population of generators, and the incidence of valid models was stored (Supplementary Figure 12). The generator with the highest incidence of valid models across all replicates was chosen for further studies.

### Neural network architecture

In REKINDLE, the GANs were implemented as conditional GANs^43^. The discriminator network was composed of three layers that have a total of 18,319 parameters: layer 1, Dense with 32 units, Dropout (0.5); layer 2, Dense with 64 units, Dropout (0.5); layer 3, Dense with 128 units, Dropout (0.5). The generator network was composed of three layers that have a total of 315,779 parameters: layer 1, Dense with 128 units, BatchNormalization, Dropout (0.5); layer 2, Dense with 256 units, BatchNormalization, Dropout (0.5); layer 3, Dense with 512 units, BatchNormalization, Dropout (0.5). We used the binary cross-entropy loss and the Adam optimizer with a learning rate of 0.0002 for training both the networks.

In RENAISSANCE, the generator neural networks were composed of three layers with 1,076,352 parameters: layer 1, dense, with 256 units, dropout (0.5); layer 2, dense, with 512 units, dropout (0.5); and layer 3, dense, with 1,024 units, dropout (0.5).

All software programs were implemented in Python (v3.6).

### Training generators to produce dynamically valid S. *cerevisiae* models

The generator for S. *cerevisiae* metabolism was constructed using the RENAISSANCE framework. The training procedure closely followed the protocol used for anaerobic E. *coli*, with minimal modifications. Firstly, we changed the allowed bounds on sampled Michaelis-Menten constants (K_M_s) to reflect physiologically relevant ranges for yeast metabolism. Specifically, K_M_ values were constrained to lie between 1.5 ∗ 10^−8^ 𝑀 and 7.4 𝑀 . Second, a gentler learning rate decay was employed, decreasing the decay factor to 𝑑 = 1%, allowing for smoother convergence during training. The neural network architecture was also expanded to accommodate the larger number of K_M_s generated in the output layer. The three fully connected layers have 1,024, 1,024, and 2,048 units in the first, second, and third layers, respectively. Training was performed over multiple iterations, and at each iteration, 100 kinetic parameter sets were generated and evaluated for dynamic validity.

To assess the flexibility and robustness of the proposed workflow across generators of varying quality, we deliberately selected generators from different stages of the training process. These generators were chosen to follow an approximately linear increase in the incidence of dynamically valid models, corresponding to early-, intermediary-, and late-stage training. Specifically, we selected a generator from iteration 40, representing an early-stage model with a low incidence of valid parameter sets (5%), a generator from iteration 62, corresponding to an intermediate stage of training with a moderate incidence (15%), and a generator from iteration 94, representing a late-stage model with a higher incidence of valid models (23%) (Figure 5A, green circles). Together, these generators span a broad range of performance and provide a systematic means of evaluating the workflow’s capability at different levels of generator maturity.

### Bioreactor simulations of anaerobically grown *E. coli*

The developed and parameterized kinetic models of anaerobic *E. coli* can be used to simulate a batch fermentation process. The bioreactor is modeled by adding two additional types of ordinary differential equations (ODEs)^53^: one for monitoring the number of cells growing in the bioreactor according to the growth rate and another that accounts for the uptake and secretion of metabolites shared between the medium and intracellular metabolism (e.g., glucose uptake and acetate secretion). We initialize the bioreactor model with steady-state concentrations and fluxes used in the kinetic model parameterization process, as well as the experimental-derived inoculum of the cells and the concentrations of the medium derived from the Varma and Palsson study.^51^ By integrating the model over 12 hours, we can compare its results with experimental fermentation data.^51^

### Non-linear integration of parameterized models

To observe the dynamic behavior of parameterized model, we randomly perturb the reference steady state metabolite concentration (𝑋_𝑟𝑒𝑓_) and flux profile of the model up to ±50%. We next integrate the ODEs using this perturbed state 𝑋’ as the initial condition 𝑋(𝑡 = 0) = 𝑋_0_. To check if a model came back to the reference steady state, we monitor the L_2_ norm of the metabolite concentrations at a given point of time 𝑋(𝑡) and the reference concentration 𝑋_0_. If the metabolite concentration at 21 mins (doubling time of *E. coli*) is less than 5% of the reference steady state we classify the model as having returned back to the steady state, i.e., we test,

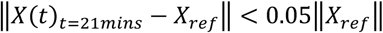

### Spearman Correlation Coefficient

For two random variables 𝑋 and 𝑌, we compute the Spearman correlation coefficient as,

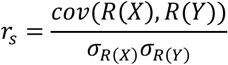

where 𝑅(𝑋) and 𝑅(𝑌) are the ranks of 𝑋 and 𝑌, respectively, 𝑐𝑜𝑣(𝑅(𝑋), 𝑅(𝑌)) the covariance of the rank variables, and 𝜎_𝑅(𝑋)_ and 𝜎_𝑅(N)_ are the standard deviations of the rank variables.

### Average deviation of simulated bioreactor responses from experimental data

The average deviation, 𝐸, between experimental data, 𝑦_𝑒𝑥𝑝_(𝑡_𝑖_), and simulated data, 𝑦_𝑒𝑥𝑝_(𝑡_𝑖_), at time points at 𝑡_𝑖_ for 𝑖 = 1, …, 𝑁, is calculated as,

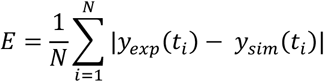

This metric was used for comparison with Varma et. al^51^, where it was first introduced.

### Jensen-Shannon (JS) Distance

For two separate probability distributions P(x) and Q(x) over the same random variable x, we can measure how different or similar these distributions are using the Jensen-Shannon divergence or distance as,

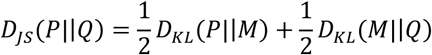

where

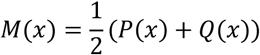

and 𝐷_𝐾𝐿_(𝑃||𝑄) is the Kullback-Leibler divergence from Q(x) to P(x) formulated as,

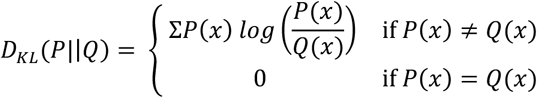

The JS divergence is symmetric and is 0 for identical distributions, with a higher divergence score representing higher dissimilarity.

### Clustering of heatmaps

The clustering of all heatmap plots in this study (Figure 2a, Supplementary Figure 1) was done using the clustermap function of the Seaborn library^67^ in Python.

## ACKNOWLEDGMENTS

This work was supported by funding from the Swiss National Science Foundation grant 200021_188623, the European Union’s Horizon 2020 research and innovation programme under grant agreement 814408, Swedish Research Council Vetenskapsradet grant 2016-06160, and the Ecole Polytechnique Fédérale de Lausanne (EPFL).

## DATA AVAILABILITY

The data supporting this study’s findings are publicly available in the Zenodo repository (https://doi.org/10.5281/zenodo.15131706 and the links therein).

## CODE AVAILABILITY

The code needed to reproduce this study’s results is publicly available at https://github.com/realLCSB/LaTeX. Python implementations of the RENAISSANCE workflow are accessible via https://github.com/EPFL-LCSB/renaissance, while the ReKinDLe workflow is available at https://github.com/EPFL-LCSB/rekindle and https://gitlab.com/EPFL-LCSB/rekindle. The ORACLE framework is implemented within the SKimPy (Symbolic Kinetic models in Python)^53^, which can be accessed at https://github.com/EPFL-LCSB/skimpy.

## CONTRIBUTIONS

S.C. and L.M. conceptualized the overall methodology, V.H. and L.M. supervised the research and acquired funding, S.C. and I.T. curated the data, models, and computational resources, S.C., O.G., J.S.B., and I.T. designed the code, S.C., I.T., O.G., and J.S.B. analyzed the data and validated results, S.C., I.T., J.S.B., and L.M. visualized the results, S.C. prepared the original draft of the manuscript, S.C., I.T., and L.M. reviewed and edited the draft. All authors read and commented on the manuscript.

## ABBREVIATIONS

ML: Machine Learning
RENAISSANCE: REconstruction of dyNAmIc models through Stratified Sampling using Artificial Neural networks and Concept of Evolution strategies
REKINDLE: REconstruction of KINetic models of metabolism using Deep Learning
TFA: Thermodynamics-based Flux Balance Analysis
ODE: Ordinary Differential Equations.

## DECLARATION OF GENERATIVE AI AND AI-ASSISTED TECHNOLOGIES IN THE WRITING PROCESS

During the preparation of this work the authors used Grammarly and ChatGPT (GPT-4) in order to improve language and text flow. After using these services, the authors reviewed and edited the content as needed and take full responsibility for the content of the publication.

## REFERENCES

1. Bordbar, A. et al. Personalized Whole-Cell Kinetic Models of Metabolism for Discovery in Genomics and Pharmacodynamics. Cell Syst. 1, 283–292 (2015).

2. Zaunseder, E. et al. Personalized metabolic whole-body models for newborns and infants predict growth and biomarkers of inherited metabolic diseases. Cell Metab. 36, 1882–1897.e7 (2024).

3. DeBerardinis, R. J. & Keshari, K. R. Metabolic analysis as a driver for discovery, diagnosis, and therapy. Cell 185, 2678–2689 (2022).

4. Yilmaz, S., Nyerges, A., van der Oost, J., Church, G. M. & Claassens, N. J. Towards next-generation cell factories by rational genome-scale engineering. Nat. Catal. 5, 751–765 (2022).

5. Lu, H., Xiao, L., Liao, W., Yan, X. & Nielsen, J. Cell factory design with advanced metabolic modelling empowered by artificial intelligence. Metab. Eng. 85, 61–72 (2024).

6. Medlock, G. L. et al. Inferring Metabolic Mechanisms of Interaction within a Defined Gut Microbiota. Cell Syst. 7, 245–257.e7 (2018).

7. Schäfer, M. et al. Metabolic interaction models recapitulate leaf microbiota ecology. Science 381, eadf5121 (2023).

8. Rios Garza, D., Gonze, D., Zafeiropoulos, H., Liu, B. & Faust, K. Metabolic models of human gut microbiota: Advances and challenges. Cell Syst. 14, 109–121 (2023).

9. Covert, M. W., Gillies, T. E., Kudo, T. & Agmon, E. A forecast for large-scale, predictive biology: Lessons from meteorology. Cell Syst. 12, 488–496 (2021).

10. Toumpe, I., Choudhury, S., Hatzimanikatis, V. & Miskovic, L. The Dawn of High-Throughput and Genome-Scale Kinetic Modeling: Recent Advances and Future Directions. ACS Synth. Biol. 10.1021/acssynbio.4c00868 (2025) doi:10.1021/acssynbio.4c00868.

11. Gelbach, P. E. et al. Kinetic and data-driven modeling of pancreatic β-cell central carbon metabolism and insulin secretion. PLOS Comput. Biol. 18, e1010555 (2022).

12. Marín-Hernández, Á. & Saavedra, E. Metabolic control analysis as a strategy to identify therapeutic targets, the case of cancer glycolysis. Biosystems 231, 104986 (2023).

13. Foster, C. et al. Assessing the impact of substrate-level enzyme regulations limiting ethanol titer in Clostridium thermocellum using a core kinetic model. Metab. Eng. 69, 286–301 (2022).

14. Narayanan, B., Weilandt, D., Masid, M., Miskovic, L. & Hatzimanikatis, V. Rational strain design with minimal phenotype perturbation. Nat. Commun. 15, 723 (2024).

15. Narayanan, B. et al. Kinetic-model-guided engineering of multiple *S. cerevisiae* strains improves *p*-coumaric acid production. Metab. Eng. 91, 430–441 (2025).

16. Zahed, M. A. et al. Kinetic modeling and half life study on bioremediation of crude oil dispersed by Corexit 9500. J. Hazard. Mater. 185, 1027–1031 (2011).

17. Kumar, M., Ji, B., Zengler, K. & Nielsen, J. Modelling approaches for studying the microbiome. Nat. Microbiol. 4, 1253–1267 (2019).

18. Gopalakrishnan, S., Dash, S. & Maranas, C. K-FIT: An accelerated kinetic parameterization algorithm using steady-state fluxomic data. Metab. Eng. 61, 197–205 (2020).

19. Choudhury, S., Narayanan, B., Moret, M., Hatzimanikatis, V. & Miskovic, L. Generative machine learning produces kinetic models that accurately characterize intracellular metabolic states. Nat. Catal. 1–13 (2024) doi:10.1038/s41929-024-01220-6.

20. Liebermeister, W. & Klipp, E. Biochemical networks with uncertain parameters. IEE Proc. - Syst. Biol. 152, 97 (2005).

21. Srinivasan, S., Cluett, W. R. & Mahadevan, R. Constructing kinetic models of metabolism at genome-scales: A review. Biotechnol. J. 10, 1345–1359 (2015).

22. Miskovic, L., Tokic, M., Fengos, G. & Hatzimanikatis, V. Rites of passage: requirements and standards for building kinetic models of metabolic phenotypes. Curr. Opin. Biotechnol. 36, 146– 153 (2015).

23. Wang, L., Birol, I. & Hatzimanikatis, V. Metabolic Control Analysis under Uncertainty: Framework Development and Case Studies. Biophys. J. 87, 3750–3763 (2004).

24. Miskovic, L. & Hatzimanikatis, V. Production of biofuels and biochemicals: in need of an ORACLE. Trends Biotechnol. 28, 391–397 (2010).

25. Mišković, L. & Hatzimanikatis, V. Modeling of uncertainties in biochemical reactions. Biotechnol. Bioeng. 108, 413–423 (2011).

26. Tran, L. M., Rizk, M. L. & Liao, J. C. Ensemble Modeling of Metabolic Networks. Biophys. J. 95, 5606–5617 (2008).

27. Haiman, Z. B., Zielinski, D. C., Koike, Y., Yurkovich, J. T. & Palsson, B. O. MASSpy: Building, simulating, and visualizing dynamic biological models in Python using mass action kinetics. PLOS Comput. Biol. 17, e1008208 (2021).

28. Andreozzi, S., Miskovic, L. & Hatzimanikatis, V. iSCHRUNK--In Silico Approach to Characterization and Reduction of Uncertainty in the Kinetic Models of Genome-scale Metabolic Networks. Metab. Eng. 33, 158–168 (2016).

29. Miskovic, L., Beal, J., Moret, M. & Hatzimanikatis, V. Uncertainty reduction in biochemical kinetic models: Enforcing desired model properties. PLoS Comput. Biol. 15, e1007242 (2019).

30. Tokic, M., Hatzimanikatis, V. & Miskovic, L. Large-scale kinetic metabolic models of Pseudomonas putida KT2440 for consistent design of metabolic engineering strategies. Biotechnol. BIOFUELS 13, 33 (2020).

31. Choudhury, S. et al. Reconstructing Kinetic Models for Dynamical Studies of Metabolism using Generative Adversarial Networks. *Nat*. Mach. Intell. 4, 710–719 (2022).

32. Massonis, G., Villaverde, A. F. & Banga, J. R. Distilling identifiable and interpretable dynamic models from biological data. PLOS Comput. Biol. 19, e1011014 (2023).

33. Weiss, K., Khoshgoftaar, T. M. & Wang, D. A survey of transfer learning. J. Big Data 3, 9 (2016).

34. Moret, M., Friedrich, L., Grisoni, F., Merk, D. & Schneider, G. Generative molecular design in low data regimes. Nat. Mach. Intell. 2, 171–180 (2020).

35. Kingma, D. P. & Welling, M. Auto-Encoding Variational Bayes. Preprint at 10.48550/arXiv.1312.6114 (2022).

36. Goodfellow, I. et al. Generative adversarial networks. Commun. ACM 63, 139–144 (2020).

37. Shen, Y., Gu, J., Tang, X. & Zhou, B. Interpreting the Latent Space of GANs for Semantic Face Editing. in 9243–9252 (2020).

38. Donahue, C., Lipton, Z. C., Balsubramani, A. & McAuley, J. Semantically Decomposing the Latent Spaces of Generative Adversarial Networks. Preprint at 10.48550/arXiv.1705.07904 (2018).

39. Bojanowski, P., Joulin, A., Lopez-Paz, D. & Szlam, A. Optimizing the Latent Space of Generative Networks. Preprint at 10.48550/arXiv.1707.05776 (2019).

40. Radford, A., Metz, L. & Chintala, S. Unsupervised Representation Learning with Deep Convolutional Generative Adversarial Networks. *arXiv.org* https://arxiv.org/abs/1511.06434v2 (2015).

41. Winant, D., Schreurs, J. & Suykens, J. A. K. Latent Space Exploration Using Generative Kernel PCA. in Artificial Intelligence and Machine Learning (eds Bogaerts, B. et al.) vol. 1196 70–82 (Springer International Publishing, Cham, 2020).

42. Samuel, D., Ben-Ari, R., Darshan, N., Maron, H. & Chechik, G. Norm-guided latent space exploration for text-to-image generation. Adv. Neural Inf. Process. Syst. 36, 57863–57875 (2023).

43. Mirza, M. & Osindero, S. Conditional generative adversarial nets. ArXiv Prepr. ArXiv14111784 (2014).

44. Shamir, M., Bar-On, Y., Phillips, R. & Milo, R. SnapShot: Timescales in Cell Biology. Cell 164, 1302–1302.e1 (2016).

45. Akbari, A., Haiman, Z. B. & Palsson, B. O. A data-driven approach for timescale decomposition of biochemical reaction networks. mSystems 9, e01001–23 (2024).

46. Saltelli, A. Making best use of model evaluations to compute sensitivity indices. Comput. Phys. Commun. 145, 280–297 (2002).

47. Saltelli, A. et al. Variance based sensitivity analysis of model output. Design and estimator for the total sensitivity index. Comput. Phys. Commun. 181, 259–270 (2010).

48. Gutenkunst, R. N. et al. Universally Sloppy Parameter Sensitivities in Systems Biology Models. PLOS Comput. Biol. 3, e189 (2007).

49. Henry, C. S., Broadbelt, L. J. & Hatzimanikatis, V. Thermodynamics-based metabolic flux analysis. Biophys. J. 92, 1792–1805 (2007).

50. Clomburg, J. M., Crumbley, A. M. & Gonzalez, R. Industrial biomanufacturing: The future of chemical production. Science 355, aag0804 (2017).

51. Varma, A. & Palsson, B. O. Stoichiometric flux balance models quantitatively predict growth and metabolic by-product secretion in wild-type Escherichia coli W3110. Appl. Environ. Microbiol. 60, 3724–3731 (1994).

52. Varma, A. & Palsson, B. O. Metabolic capabilities of Escherichia coli: I. Synthesis of biosynthetic precursors and cofactors. J. Theor. Biol. 165, 477–502 (1993).

53. Weilandt, D. R. et al. Symbolic kinetic models in python (SKiMpy): intuitive modeling of large-scale biological kinetic models. Bioinformatics 39, btac787 (2023).

54. Salvy, P. et al. pyTFA and matTFA: a Python package and a Matlab toolbox for Thermodynamics-based Flux Analysis. Bioinforma. Oxf. Engl. 35, 167–169 (2019).

55. Ataman, M., Hernandez Gardiol, D. F., Fengos, G. & Hatzimanikatis, V. redGEM: Systematic reduction and analysis of genome-scale metabolic reconstructions for development of consistent core metabolic models. PLoS Comput. Biol. 13, e1005444 (2017).

56. Ataman, M. & Hatzimanikatis, V. lumpGEM: Systematic generation of subnetworks and elementally balanced lumped reactions for the biosynthesis of target metabolites. PLoS Comput. Biol. 13, e1005513 (2017).

57. Mavrovouniotis, M. L. Group contributions for estimating standard gibbs energies of formation of biochemical compounds in aqueous solution. Biotechnol. Bioeng. 36, 1070–1082 (1990).

58. Jankowski, M. D., Henry, C. S., Broadbelt, L. J. & Hatzimanikatis, V. Group contribution method for thermodynamic analysis of complex metabolic networks. Biophys. J. 95, 1487–1499 (2008).

59. Liebermeister, W. & Klipp, E. Bringing metabolic networks to life: convenience rate law and thermodynamic constraints. Theor. Biol. Med. Model. 3, 41 (2006).

60. Hanekom, A. J. Generic kinetic equations for modelling multisubstrate reactions in computational systems biology /.

61. Chang, A. et al. BRENDA, the ELIXIR core data resource in 2021: new developments and updates. Nucleic Acids Res. 49, D498–D508 (2021).

62. Gibson, B., Wilson, D. J., Feil, E. & Eyre-Walker, A. The distribution of bacterial doubling times in the wild. Proc. R. Soc. B Biol. Sci. 285, 20180789 (2018).

63. Salimans, T., Ho, J., Chen, X., Sidor, S. & Sutskever, I. Evolution Strategies as a Scalable Alternative to Reinforcement Learning. 10.48550/ARXIV.1703.03864 (2017) doi:10.48550/ARXIV.1703.03864.

64. Vent, W. Rechenberg, Ingo, Evolutionsstrategie — Optimierung technischer Systeme nach Prinzipien der biologischen Evolution. 170 S. mit 36 Abb. Frommann-Holzboog-Verlag. Stuttgart 1973. Broschiert. Feddes Repert. 86, 337–337 (1975).

65. Soh, K. C., Miskovic, L. & Hatzimanikatis, V. From network models to network responses: integration of thermodynamic and kinetic properties of yeast genome-scale metabolic networks. FEMS Yeast Res. 12, 129–143 (2012).

66. Chakrabarti, A., Miskovic, L., Soh, K. C. & Hatzimanikatis, V. Towards kinetic modeling of genome-scale metabolic networks without sacrificing stoichiometric, thermodynamic and physiological constraints. Biotechnol. J. 8, 1043–1057 (2013).

67. Waskom, M. seaborn: statistical data visualization. J. Open Source Softw. 6, 3021 (2021).

