## Supplementary Material for "Generative Approaches to Kinetic Parameter Inference in Metabolic Networks via Latent Space Exploration"

##### Abbreviations:

ACALD: Acetaldehyde dehydrogenase (acetylating), ACALDpp: Acetaldehyde dehydrogenase (periplasmic), AC2rpp: Acetate transport via proton symporter (reversible, periplasmic), ACKr: Acetate kinase (reversible), ACONIt: Aconitase, ACS: Acetyl-CoA synthetase, AKGDH: Alpha-ketoglutarate dehydrogenase complex, ALCD2x: Alcohol dehydrogenase (cytoplasmic), ATP4rpp: ATP synthase/transport reaction (periplasmic), Biomass: Biomass objective function, CO2tpp: $CO_2$ transport (periplasmic), CS: Citrate synthase, CYTBD3: Cytochrome bd oxidase, E3: Dihydrolipoamide dehydrogenase (E3 component), ETOHpp: Ethanol transport (periplasmic), FBA: Fructose-bisphosphate aldolase, FBP: Fructose-1,6-bisphosphatase, F6P: Fructose 6-phosphate, G6PD: Glucose-6-phosphate 1-dehydrogenase, GAPD: Glyceraldehyde-3-phosphate dehydrogenase, GDH: Glutamate dehydrogenase, GOGAT: Glutamine oxoglutarate aminotransferase, ICL: Isocitrate lyase, MALS: Malate synthase, MALt: Malate transport, MDH: Malate dehydrogenase, ME1: Malic enzyme ($NAD^+$-dependent), NADH2: NADH:ubiquinone oxidoreductase, NADTRHD: Transhydrogenase (NAD(P) transhydrogenase), NDH2: NADH dehydrogenase II, O2tpp: $O_2$ transport (periplasmic), PDH: Pyruvate dehydrogenase complex, PFK: Phosphofructokinase, PFK_3: Alternative phosphofructokinase isoform, PGI: Phosphoglucose isomerase, PGK: Phosphoglycerate kinase, PGM: Phosphoglycerate mutase, PGP: Phosphoglycolate phosphatase, PPC: Phosphoenolpyruvate carboxylase, PPK: Polyphosphate kinase, PTAr: Phosphotransacetylase (reversible), PYK: Pyruvate kinase, RPE: Ribulose-5-phosphate 3-epimerase, RPI: Ribose-5-phosphate isomerase, SUCDi: Succinate dehydrogenase (irreversible), SUCOAS: Succinyl-CoA synthetase, THD2pp: Transhydrogenase (periplasmic), TK1: Transketolase 1, TPI: Triose-phosphate isomerase. ru5p: D-Ribulose 5-phosphate, dhap: Dihydroxyacetone phosphate, nadph: Nicotinamide adenine dinucleotide phosphate – reduced, r5p: Alpha-D-Ribose 5-phosphate, coa: coenzyme A, oaa: Oxaloactetate

##### Supplementary Note 1: Augmenting latent space of neural networks for generating kinetic parameter sets

Generative deep learning frameworks, like REKINDLE and RENAISSANCE use, adversarial training and Natural Evolutionary Strategies respectively, to obtain generator neural networks, $G(\boldsymbol{w})$, that can consistently generate dynamically desirable kinetic parameter sets given a kinetic structure (in the form of a system of ODEs, $dx/dt = \boldsymbol{S}.\boldsymbol{v}(\boldsymbol{x}, \boldsymbol{k}, E, t)$ where $\boldsymbol{S}$ is stoichiometry of the metabolic network), steady state vector of fluxes, $\boldsymbol{v}$, and concentrations, $\boldsymbol{x}$, and a mathematically defined design objective. Both of these methods, map a simplified latent space (multivariate standard Gaussian space), $\boldsymbol{z}$, to the subspace of desired kinetic parameters, $\boldsymbol{k}$, given the model constraints (Figure 1a) as,

$$G(\boldsymbol{z} \mid \boldsymbol{w}, \boldsymbol{S}.\boldsymbol{v}, \{\boldsymbol{x}, \boldsymbol{v}\}) = \boldsymbol{k}$$

where $\boldsymbol{w}$ are the parameters of the neural network that are optimized during training. Once the optimized weights, $\boldsymbol{w}_{opt}$, are obtained, a trained neural network is used to generate new parameter sets by sampling random vectors, $\boldsymbol{z}^*$, from the latent space,

$$G(\boldsymbol{z}^* \mid \boldsymbol{w}_{opt}) = \boldsymbol{k}^*$$

Due to the nonlinearity of the generator neural network, $G(\boldsymbol{w}_{opt})$, modifying the input latent vector, $\boldsymbol{z}^*$, results in a new kinetic parameter set,

$$G(\boldsymbol{z}^* + \boldsymbol{\Delta z} \mid \boldsymbol{w}_{opt}) = \boldsymbol{k}^{new}$$

Thus, simple linear transformations in the features of the latent input vector gives rise to different kinetic parameter sets which when parameterized in the dynamic model leads to the different dynamic properties from the original model.

#### Supplementary Note 2: Co-occurrence of dynamically important enzymes for a steady state profile of fluxes and concentration.

In Figure 4, we used latent space manipulation to uncover general dynamic bottlenecks of the central carbon metabolism of *E. coli*. We also investigated whether certain kinetic parameters or enzymes occur concurrently as the important factors that determine dynamic properties, as this would reveal the role of different modalities of a metabolic network that dictate overall behaviour. For this we checked the important enzymes for each model across 4 different steady states to check if such significant co-occurrences emerged (Supplementary Figure 1). We observed that certain enzyme couples co-occur more frequently than others, but the results differ significantly across steady states and are not conclusive.

We also checked if there was any preference for enzymes or associated kinetic parameters emerging as the primary bottlenecks with respect to the $\tau_{max}$ of the model being investigated for each steady state. But again, we observed no conclusive patterns in the occurrences of important enzymes and the dominant time constant, $\tau_{max}$, of the model. Even though the results of this analysis were inconclusive, we believe that finding such enzyme co-occurrences and direct relations between enzymes and the dynamic properties of the network remain important open problems and merit thorough investigation in future studies.

#### Supplementary Note 3: Comparison between Latent Space Sensitivity Analysis and Variance-Based Global Sensitivity Analysis.

Perturbations applied in latent space must ultimately be mapped back to the kinetic parameter space for numerical integration and evaluation of system dynamics. In this sense, latent space exploration and parameter-space sensitivity analysis are not conceptually disjoint. However, a key distinction lies in how perturbations are applied. Perturbing individual latent dimensions induces *structured and coordinated* changes across all kinetic parameters, reflecting the learned low-dimensional manifold of dynamically feasible models. By contrast, traditional variance-based global sensitivity analysis (GSA) perturbs parameters individually or in explicitly enumerated combinations.
As a consequence, latent space perturbations probe collective parameter modes that are most relevant for macroscopic dynamic behavior, whereas parameter-wise GSA evaluates marginal contributions of individual parameters under assumed independence and predefined sampling ranges.

Comprehensive variance-based GSA rapidly becomes computationally prohibitive for large-scale kinetic models. The *E. coli* kinetic model used in the first three studies comprises $D = 258$ kinetic parameters. Using the Saltelli sampling scheme[1,2] with a base sample size $N$, all Sobol indices up to order $M$ can be estimated with $N(D + M)$ model evaluations, with each additional interaction order requiring only $+N$ additional simulations. In other words, the number of required model evaluations scales as: (i) 1$^{st}$-order Sobol indices: $N \times (D + 1)$; (ii) 1$^{st}$- and total-order indices: $N \times (D + 1)$, (iii) 1$^{st}$-, 2$^{nd}$-, and total order indices, $N \times (D + 2)$. Using a relatively coarse-grained estimate ($N = 256$), computing first-order Sobol indices requires 66,304 model evaluations. Increasing precision to $N = 1024$ raises this number to 265,216 evaluations. Inclusion of second-order and total-order effects further increases the computational burden to 66,560 ($N = 256$) or 266,240 ($N = 1024$) evaluations, with even higher-order interactions scaling even more unfavourably. By contrast, the latent space–based sensitivity analysis presented in Figure 2 required only 1,681 model evaluations. This represents a reduction of more than two orders of magnitude in computational cost while still capturing substantial variance in system-level outputs.

We computed 1$^{st}$-order Sobol indices for all 258 kinetic parameters of the *E. coli* model used in the first three studies. A high-precision estimate was obtained using $N = 1024$. The resulting distribution of Sobol indices is shown in Supplementary Figure 18. All 1$^{st}$-order Sobol fall within a narrow range between approximately 0.0037 and 0.004, corresponding to sensitivities of roughly 3.5–4%. No individual parameter dominates the variance of the dominant time constant $\tau_{\max}$. This behavior is consistent with prior observations that kinetic parameters in biochemical networks are typically *sloppy* with respect to macroscopic dynamics, such that large variations in individual parameters have limited and difficult-to-interpret effects on system-level behavior.

The narrow distribution of Sobol indices illustrates a key limitation of parameter-wise GSA in large kinetic models: while formally comprehensive, it often fails to identify interpretable control parameters or dominant regulatory mechanisms. In contrast, latent space exploration enables the identification of specific kinetic parameters whose *coordinated variation* strongly affects dynamic behavior, even when those parameters exhibit weak individual Sobol sensitivities.

Because latent variables correspond to directions of maximal variation in dynamically feasible model space, they provide a compact representation of effective control coordinates. Mapping influential latent directions back to parameter space therefore highlights combinations of parameters that jointly modulate system dynamics, rather than isolated parameter effects.

Variance-based GSA captures parameter interactions only through explicit computation of higher-order indices, which incurs substantial additional computational cost and scales poorly with model dimensionality. Latent space exploration, by contrast, perturbs all parameters simultaneously along learned directions and therefore *implicitly captures higher-order interactions* without requiring their explicit enumeration.

Using the same statistical framework described in the main manuscript, we analyzed parameter and enzyme co-occurrence statistics following latent perturbations (Supplementary Note 1). While no statistically significant interaction clusters were detected in the present case, the framework itself is general and enables interaction analysis without additional model evaluations. Taken together, these results indicate that latent space–based sensitivity analysis provides a scalable and computationally efficient alternative to traditional variance-based GSA for large kinetic models. By operating on a low-dimensional manifold of dynamically relevant parameter combinations, latent space exploration circumvents the combinatorial explosion associated with exhaustive parameter-wise perturbations while naturally incorporating higher-order interactions. This efficiency advantage is particularly important for near-genome-scale kinetic models, where conventional GSA rapidly becomes infeasible.

### Supplementary Figures

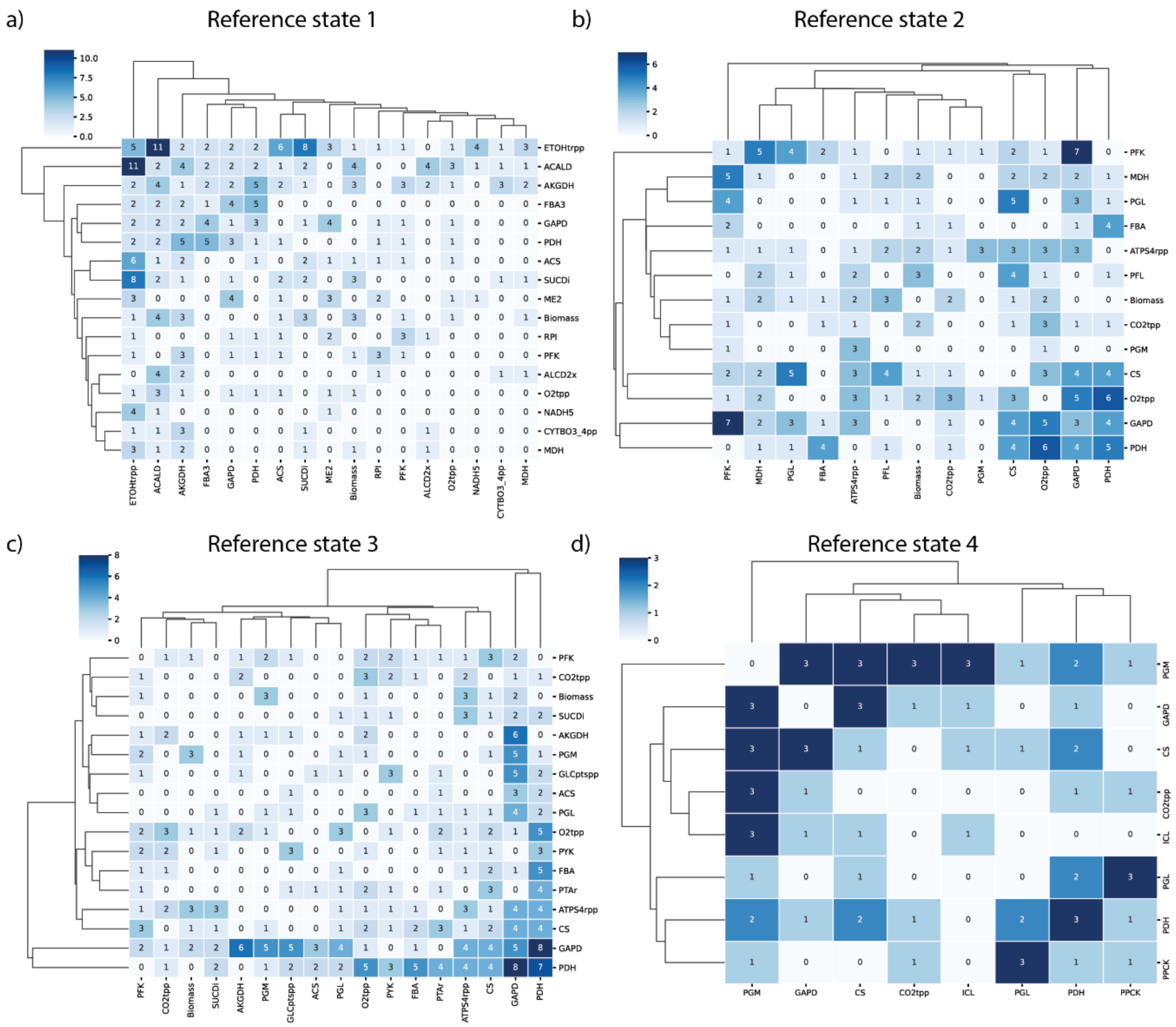


**Supplementary Figure 1**: The co-occurences of enzymes whose associated kinetic parameters emerge as the primary bottlenecks (top 3) for determining $\tau_{max}$ of the network for 4 different steady state profiles represented in (a) (b) (c) and (d) as explained in Figure 4. The color bar represents the number of co-occurences with darker shades of blue representing higher number of co-occurences.

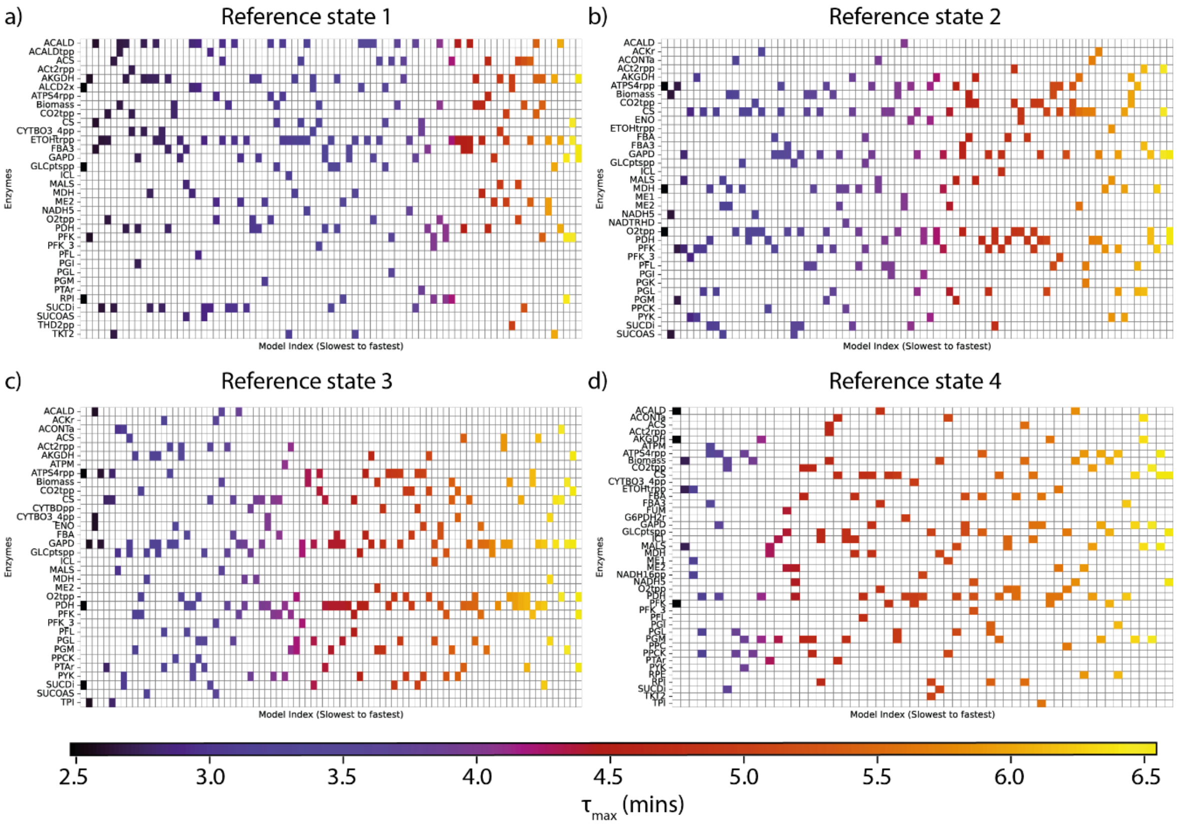


**Supplementary Figure 2**: Enzymes which whose associated kinetic parameters emerge as the top 3 parameters affecting $\tau_{max}$ of the *E. coli* central carbon metabolism network for 4 different steady state profiles. The models are sorted from slowest (dark blue) to fastest (yellow) depending on their dominant time constant $\tau_{max}$.

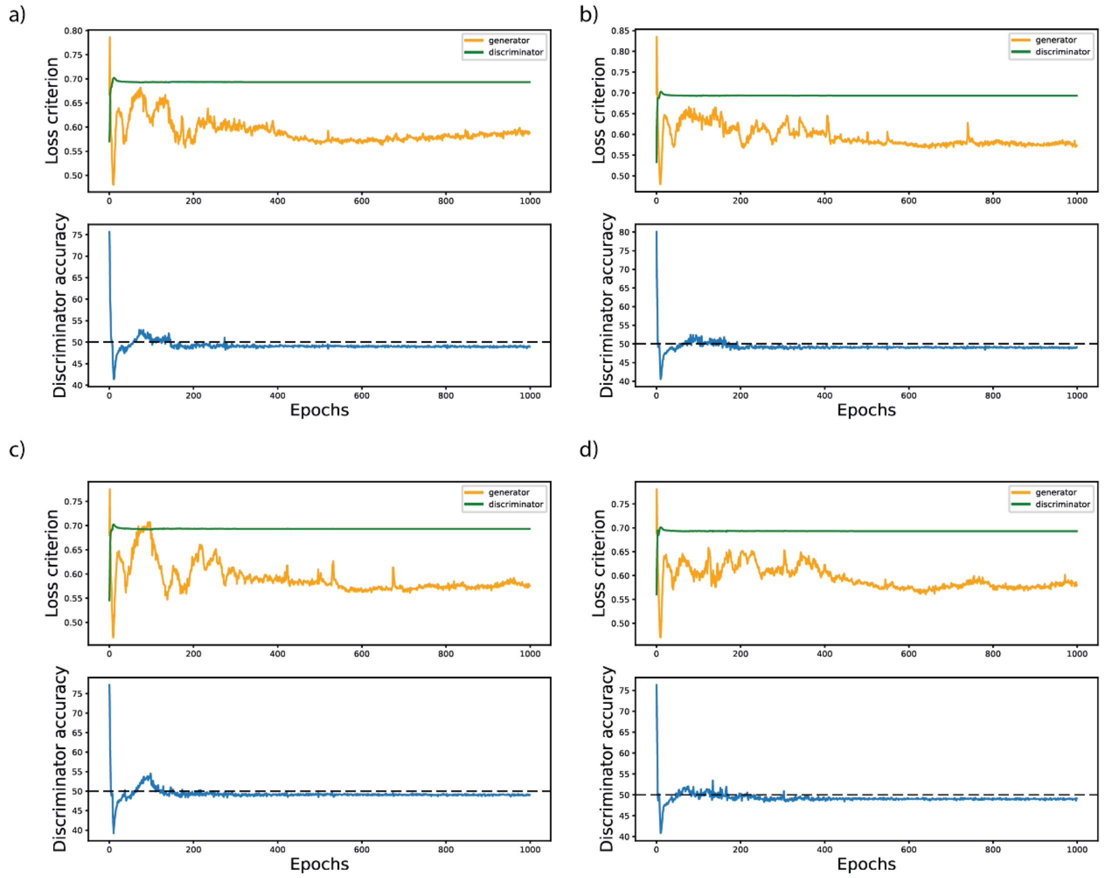


**Supplementary Figure 3**: (Upper panel) Discriminator and generator loss (Lower Panel) Discriminator accuracy during REKINDLE training for (a) steady state 1 (b) steady state 2 (c) steady state 3 (d) steady state 4 as described in the Methods section for *E. coli* central carbon metabolism.

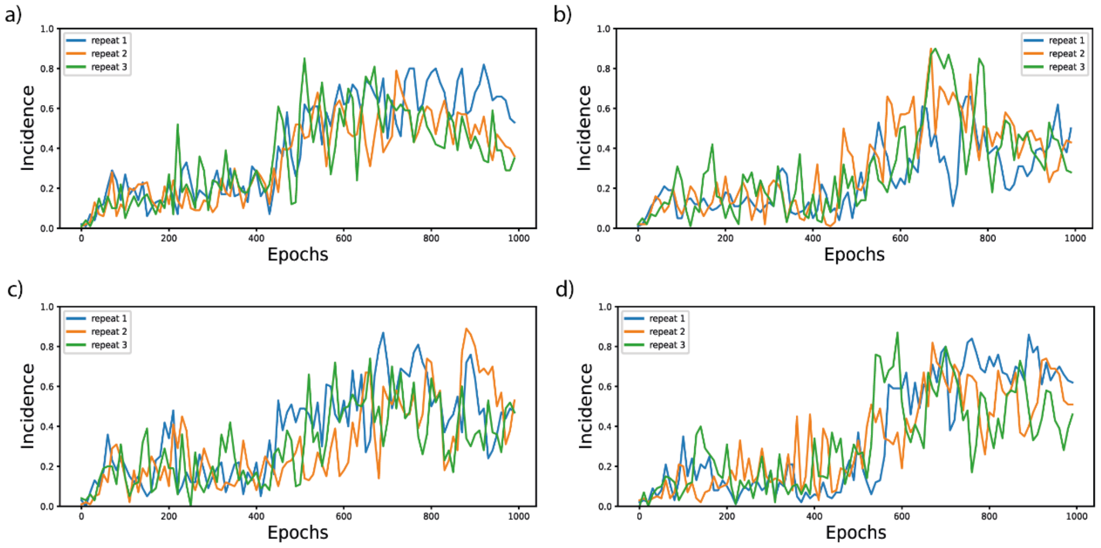


**Supplementary Figure 4**: Incidence of valid models for (a) steady state 1 (b) steady state 2 (c) steady state 3 (d) steady state 4, with 3 statistical repeats in each case as described in the Methods section for *E. coli* central carbon metabolism.

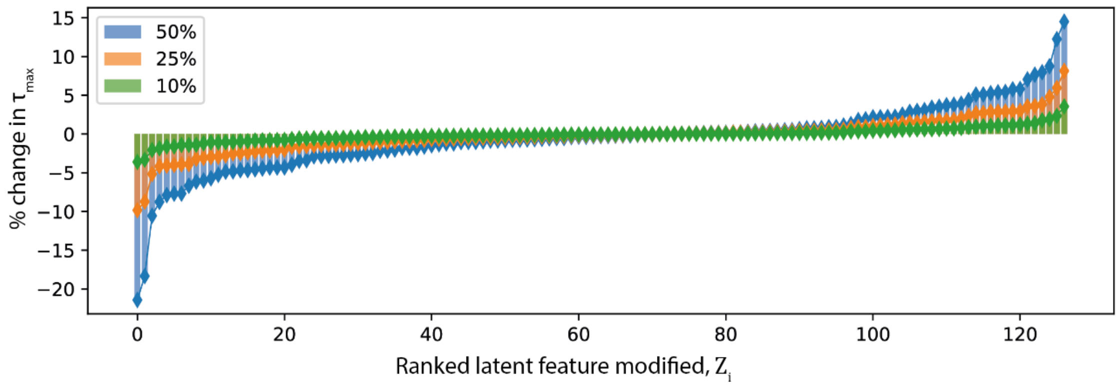


**Supplementary Figure 5:** Percentage change in dominant time constant ($\tau_{max}$) of the model analyzed in Figure 2 when each latent feature is increased by 10% (green), 25% (orange) and 50% (blue) respectively.

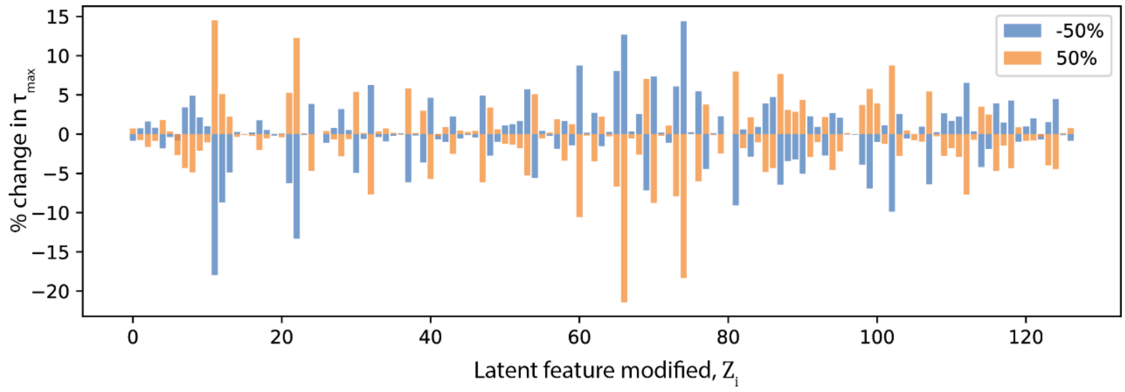


**Supplementary Figure 6:** Percentage change in dominant time constant ($\tau_{max}$) of the model analyzed in Figure 2 when each latent feature is increased by 50% (orange) and decreased by 50% (blue) respectively.

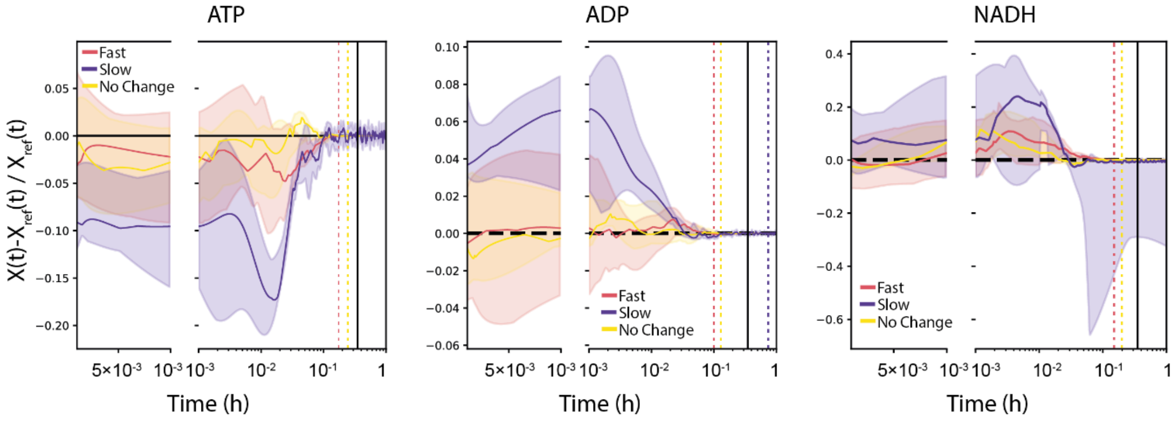


**Supplementary Figure 7:** Normalized dynamic responses of (left) ATP (middle) ADP and (right) NADP concentrations of chosen models from Figure 2c (black squares). Black line indicates doubling time of WT *E. coli*. Color dashed lines represent time points when median response comes back to within 5% of the reference steady state value. Note that for ATP and NADH the slow models (purple) do not come back to steady state in 1 hour.

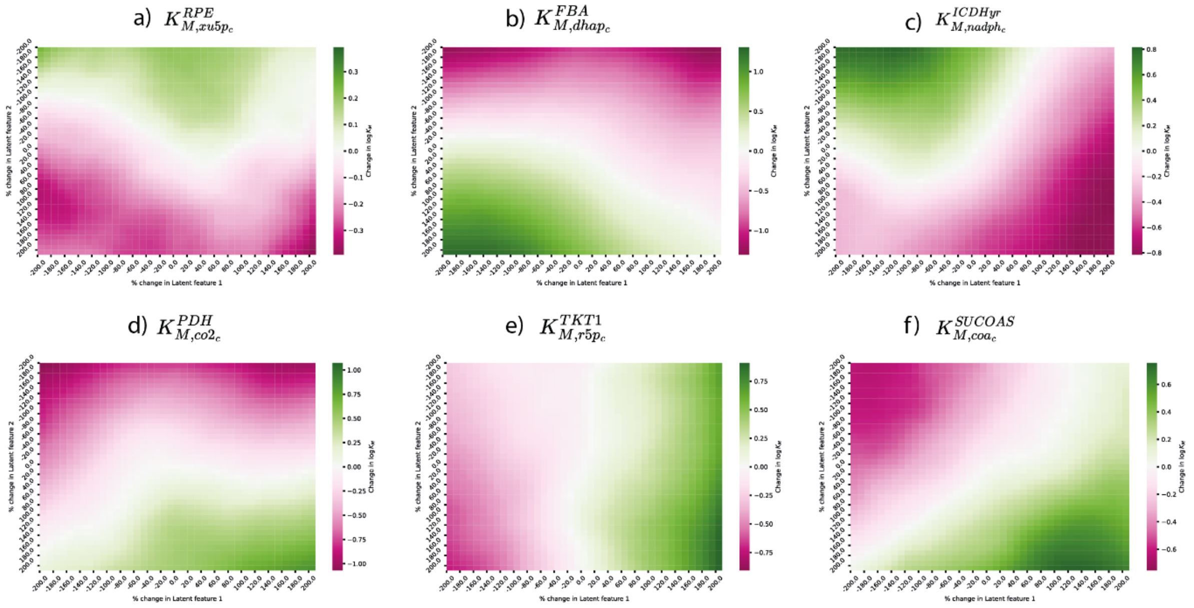


**Supplementary Figure 8:** Non exhaustive examples of change in individual kinetic parameters values for 1681 models obtained in Figure 2c for (a)$K^{RPE}_{M,xu5p}$(b) $K^{FBA}_{M,dhap}$ (c) $K^{ICDHyr}_{M,nadph}$ (d) $K^{PDH}_{M,co2}$ (e) $K^{TKT1}_{M,r5p}$ (f) $K^{SUCOAS}_{M,coa}$. For exhaustive list check supplementary data and instructions for generating figures.

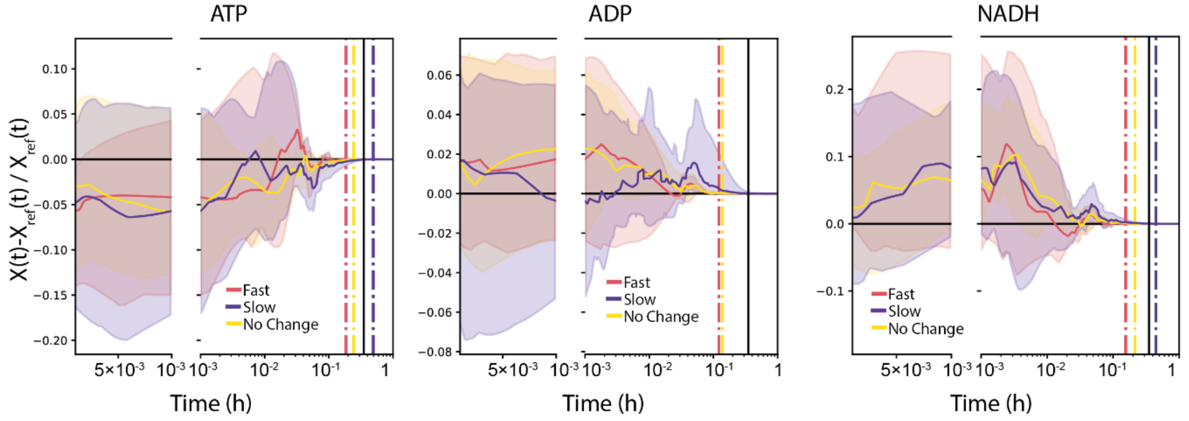


**Supplementary Figure 9:** Dynamic responses of normalized (left) ATP (middle) ADP and (right) NADP concentrations of chosen models from Figure 3c (black squares). Black line indicates doubling time of WT *E. coli*. Color dashed lines represent time points when median response comes back to within 1% of the reference steady state value. Note that for ADP the slow models (purple) do not come back to steady state in 1 hour.

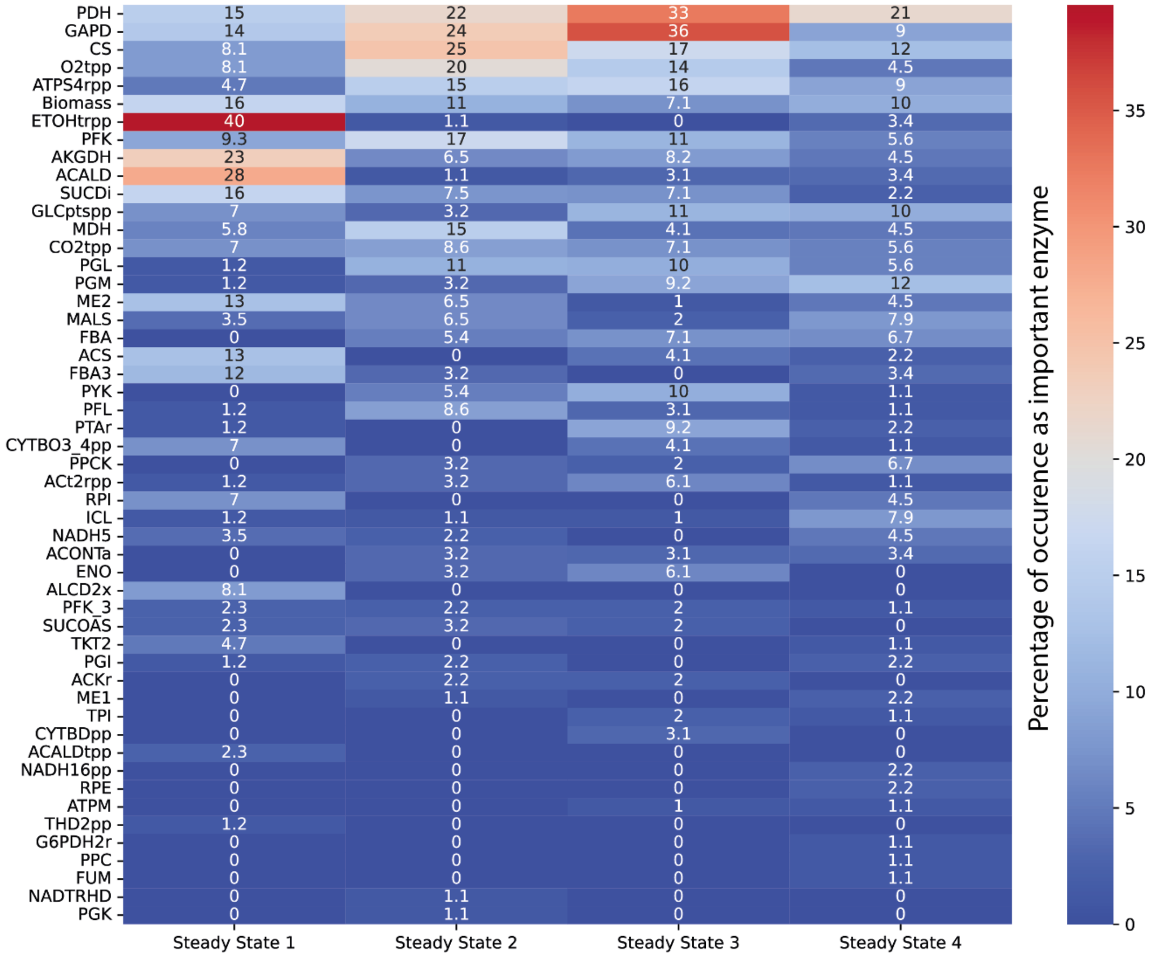


**Supplementary Figure 10:** Percentage occurrence of enzymes whose associated kinetic parameters strongly affect $\tau_{max}$ of the central carbon *E. coli* metabolism for 4 different steady state profiles. Red color denotes high occurrence and blue denotes low occurrence.

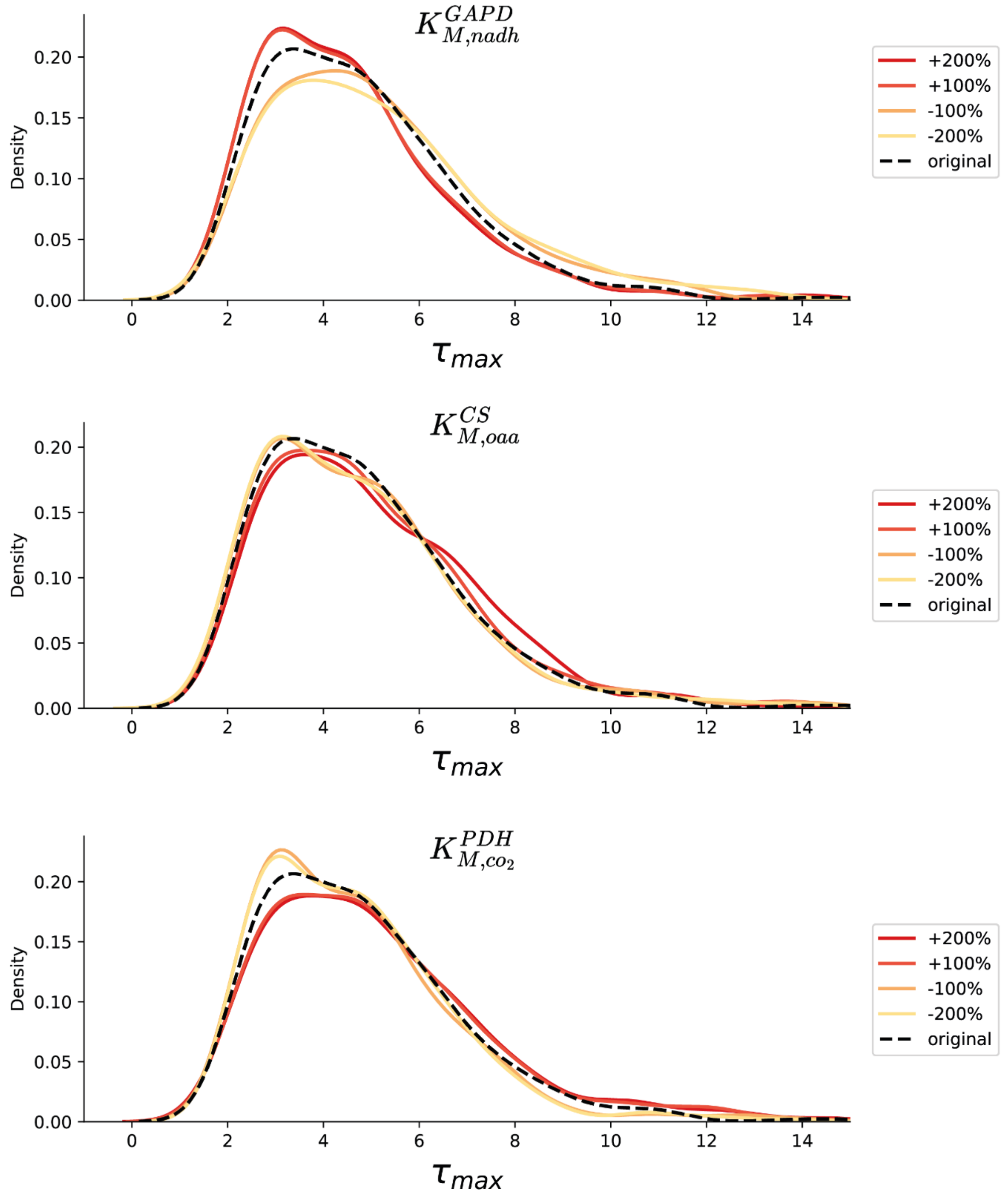


**Supplementary Figure 11:** Distribution of $\tau_{max}$ of models when (top) $K^{GAPD}_{m,nadh}$ (middle) $K^{CS}_{m,oaa}$ (bottom) $K^{PDH}_{m,co2}$ are modified from their original values by different magnitudes (colored lines). The black dashed line represents the distribution of $\tau_{max}$ of the original unperturbed models.

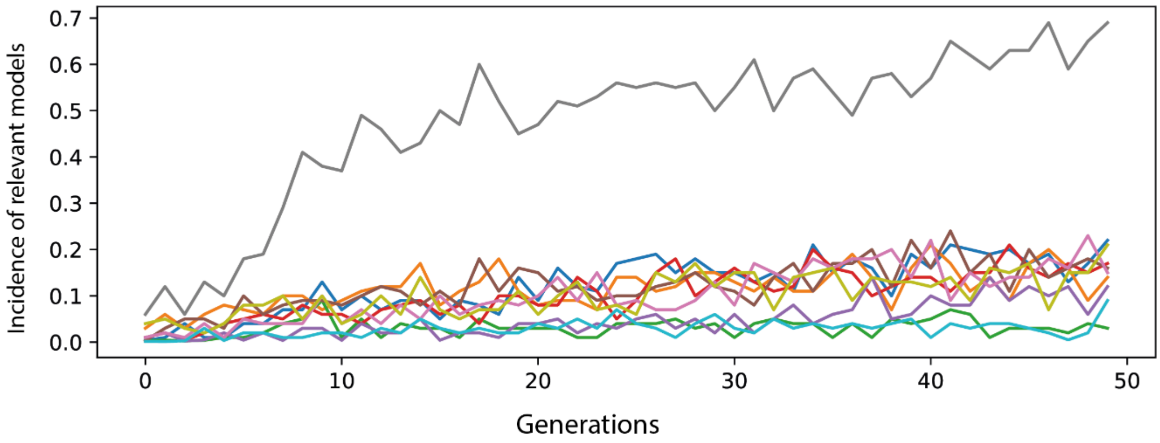


**Supplementary Figure 12:** Incidence of relevant models (Methods) for anaerobic *E. coli* metabolism obtained using RENAISSANCE for 50 generations for 10 statistical repeats (different colored lines). The repeat with the highest incidence (gray line) was selected for further studies.

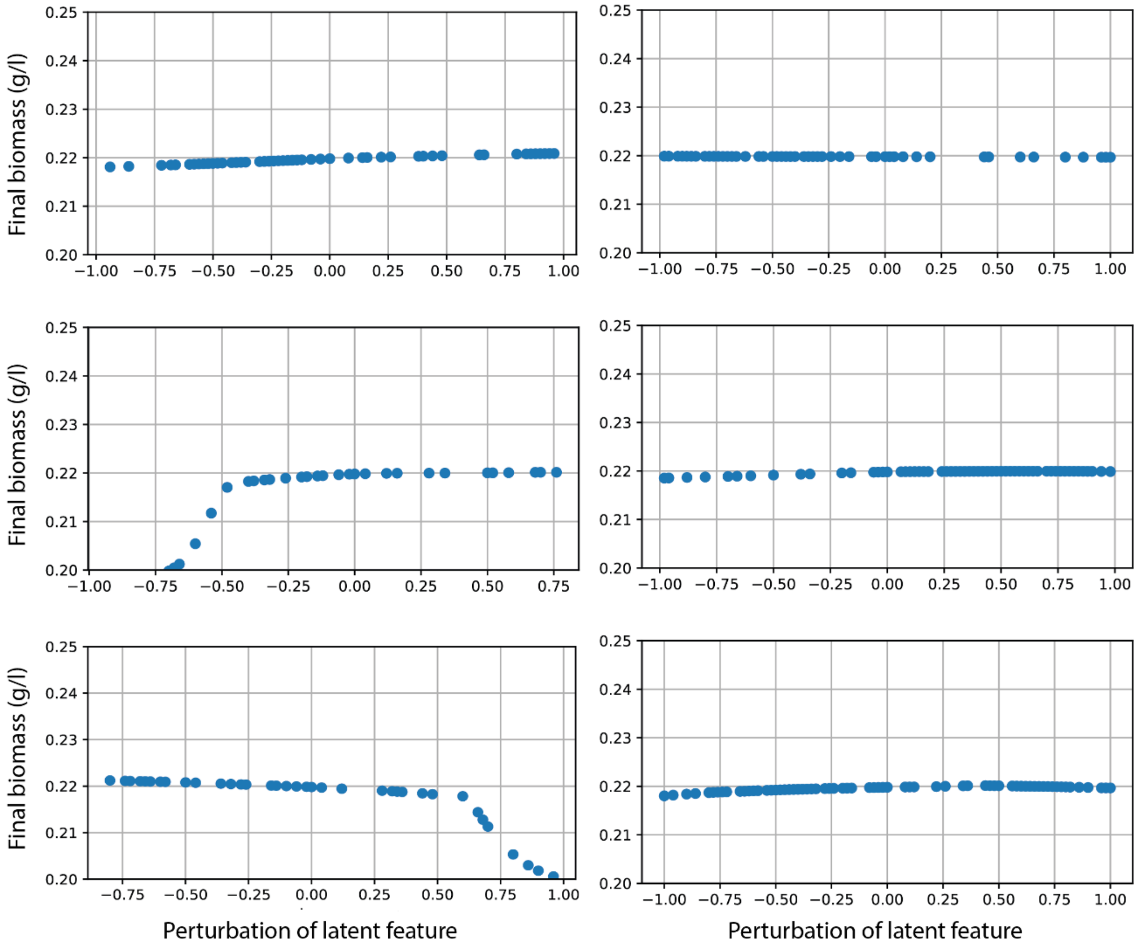


**Supplementary Figure 13:** Non exhaustive visualization of final biomass titers of model responses when individual latent features are perturbed from their original values (as explained in Figure 6). For exhaustive list check supplementary data and instructions for generating figures.

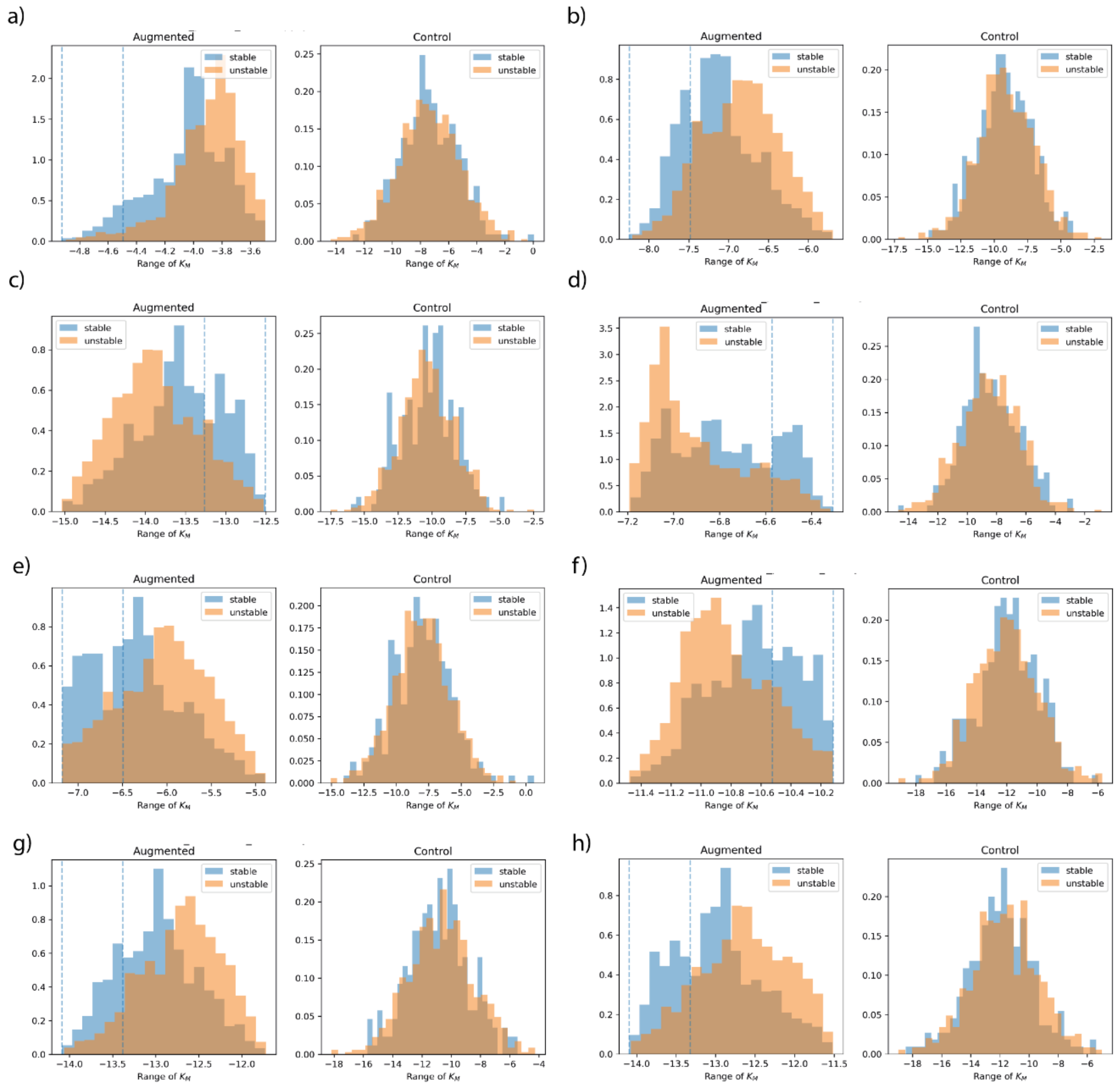


**Supplementary Figure 14:** Distributions of the parameter (a) $K_{M,lac\_D}^{D-LACt2pp}$ (b) $K_{M,ru5p\_D}^{RPI}$ (c) $K_{M,nadh}^{GAPD}$ (d) $K_{M,nadph}^{GND}$ (e) $K_{M,atp}^{GLNS}$ (b) $K_{M,pyr}^{PPS}$ (c) $K_{M,nadh}^{NADH10}$ (d) $K_{M,h}^{Biomass}$ in generated models when (left) models are obtained through systematic perturbation of the latent point of the model in Fig 6a (right) when models are obtained through random sampling of the latent space of the trained generator. Gray vertical lines in top panel represent the 30th percentile values for this parameter that favors stability.

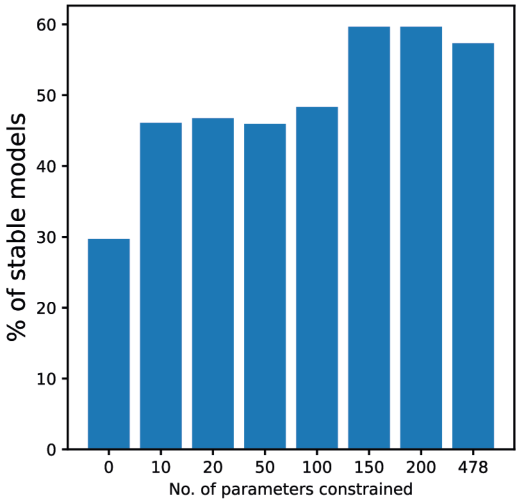


**Supplementary Figure 15:** Percentage of stable models of anaerobic *E. coli* metabolism achieved when different number of individual parameters are constrained as explained in Figure 6e and Supplementary Figure 12.

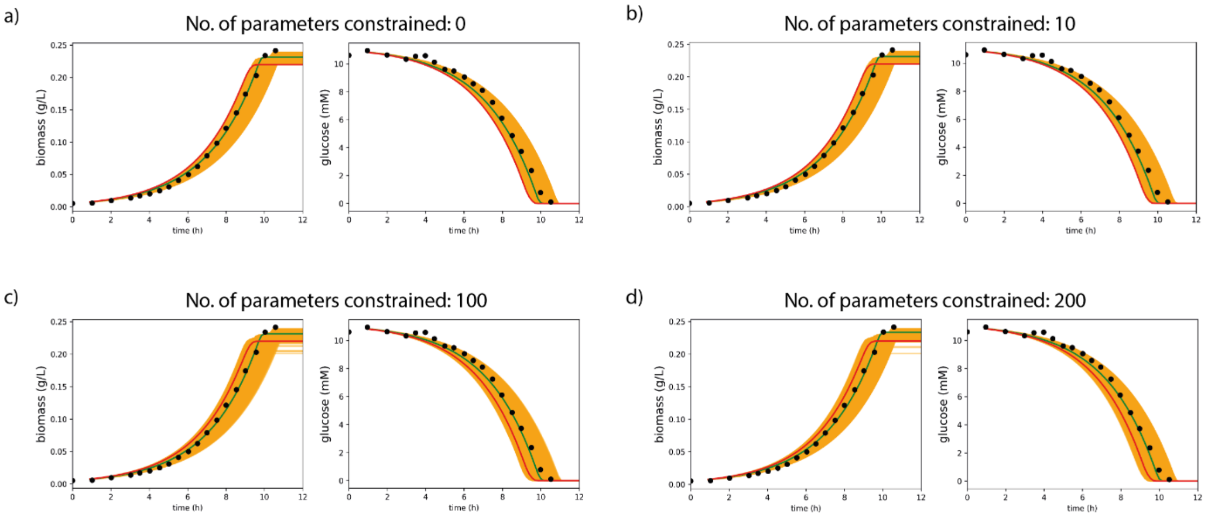


**Supplementary Figure 16:** Simulated bioreactor responses of locally stable models of anerobic *E. coli* metabolism obtained by perturbing latent features of the original model when (a) No parameters (b) 10 parameters (c) 100 parameters and (d) 200 parameters are constrained to ranges that favor stability as explained in Figure 6, Supplementary Figure 12 and Supplementary Figure 13. Red line represents the original unperturbed model and green line represents the best model obtained with lowest average deviation (Methods) from experimental values. Black dots represent experimental values.

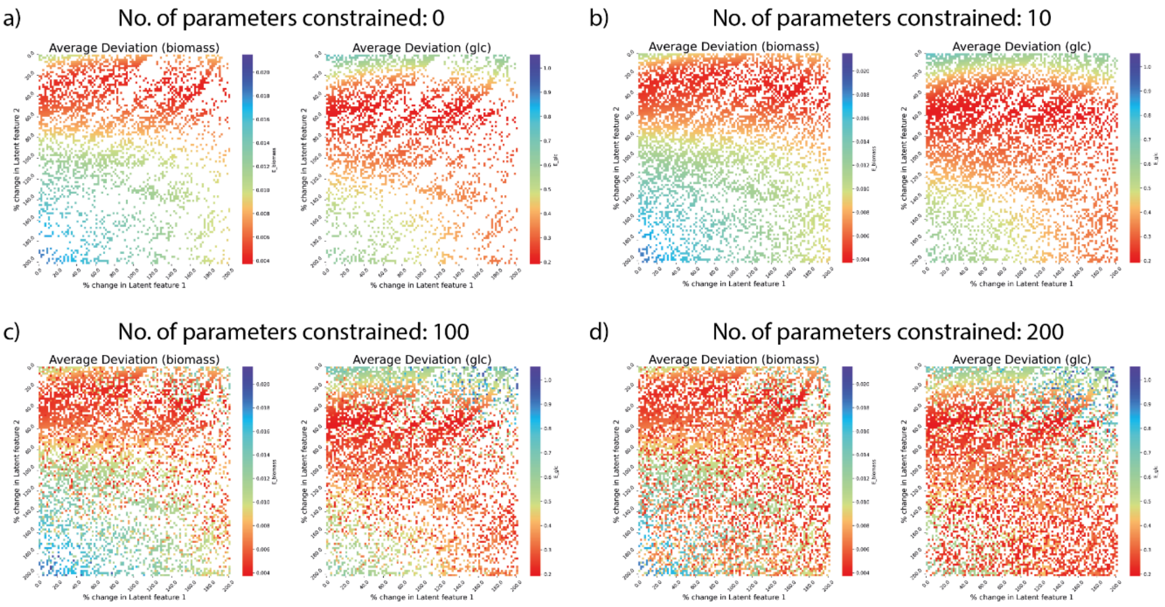


**Supplementary Figure 17:** Average deviations of simulated bioreactor responses of anaerobic *E. coli* metabolism for (left) biomass and (bottom) glucose from experimental bioreactor datapoints when the two latent features in Figure 6b are perturbed simultaneously and (a) no parameters (b) 10 parameters (c) 100 parameters and (d) 200 parameters are constrained to ranges that favor stability as explained in Figure 6, Supplementary Figure 12 and Supplementary Figure 13.

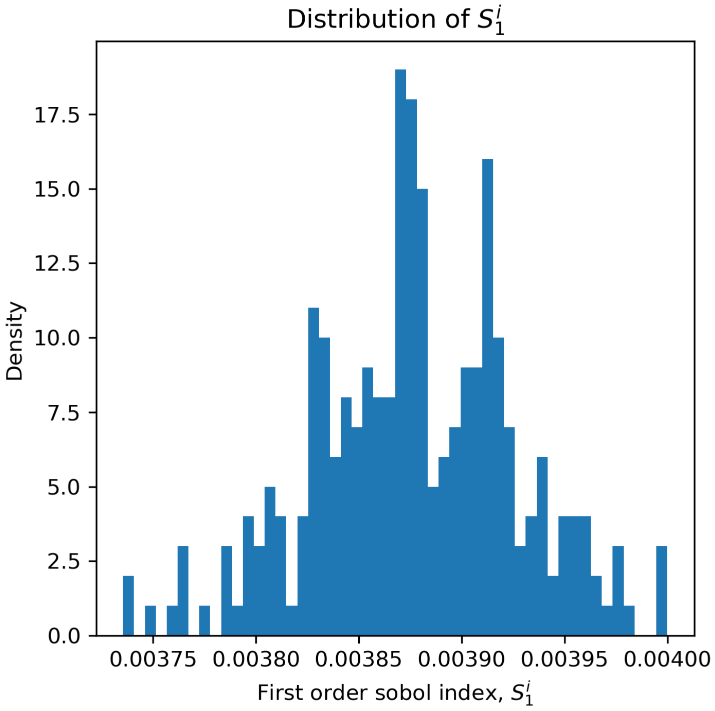


**Supplementary Figure 18:** Distribution of 1st-order Sobol indices for the *E. coli* kinetic model comprising 258 kinetic parameters.

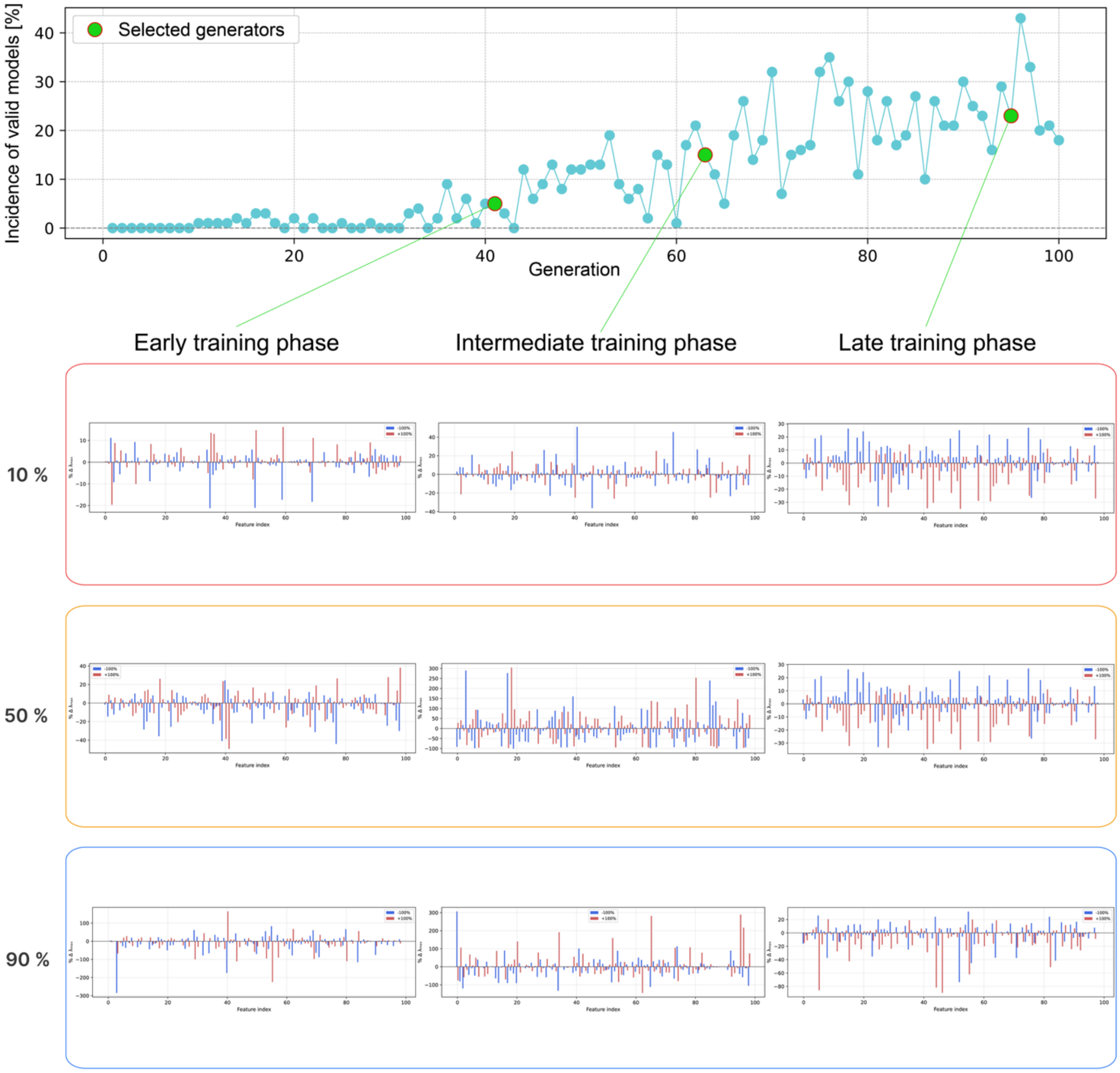


**Supplementary Figure 19:** Perturbation analysis for identifying the most effective latent features. For each generator and each latent-space sample (corresponding to the 10th, 50th, and 90th percentiles of the sampled latent distribution), we performed a ±100% perturbation of each latent feature individually and recorded the resulting percentage change in the maximum eigenvalue ($\tau_{max}$). In each case, we identified the two latent features exhibiting the largest relative difference between the −100% and +100% perturbations. For the early-stage generator, the selected latent features were: 10th percentile ($z_1$ = 50, $z_2$ = 35); 50th percentile ($z_1$ = 77, $z_2$ = 98); and 90th percentile ($z_1$ = 40, $z_2$ = 55). For the intermediate-stage generator, the selected latent features were: 10th percentile ($z_1$ = 41, $z_2$ = 73); 50th percentile ($z_1$ = 18, $z_2$ = 17); and 90th percentile ($z_1$ = 65, $z_2$ = 0). For the late-stage generator, the selected latent features were: 10th percentile ($z_1$ = 52, $z_2$ = 15); 50th percentile ($z_1$ = 75, $z_2$ = 47); and 90th percentile ($z_1$ = 5, $z_2$ = 44).

### References

1. Saltelli, A. Making best use of model evaluations to compute sensitivity indices. *Comput. Phys. Commun.* **145**, 280–297 (2002).
2. Saltelli, A. *et al.* Variance based sensitivity analysis of model output. Design and estimator for the total sensitivity index. *Comput. Phys. Commun.* **181**, 259–270 (2010).
